# Wnt/β-catenin signaling within multiple cell types dependent upon *kramer* regulates *Drosophila* intestinal stem cell proliferation

**DOI:** 10.1101/2023.02.21.529411

**Authors:** Hongyan Sun, Adnan Shami Shah, Alessandro Bonfini, Nicolas S. Buchon, Jeremy M. Baskin

## Abstract

The gut epithelium is subject to constant renewal, a process reliant upon intestinal stem cell (ISC) proliferation that is driven by Wnt/β-catenin signaling. Despite the importance of Wnt signaling within ISCs, the relevance of Wnt signaling within other gut cell types and the underlying mechanisms that modulate Wnt signaling in these contexts remain incompletely understood. Using challenge of the *Drosophila* midgut with a non-lethal enteric pathogen, we examine the cellular determinants of ISC proliferation, harnessing *kramer*, a recently identified regulator of Wnt signaling pathways, as a mechanistic tool. We find that Wnt signaling within Prospero-positive cells supports ISC proliferation and that *kramer* regulates Wnt signaling in this context by antagonizing *kelch*, a Cullin-3 E3 ligase adaptor that mediates Dishevelled polyubiquitination. This work establishes *kramer* as a physiological regulator of Wnt/β-catenin signaling in vivo and suggests enteroendocrine cells as a new cell type that regulates ISC proliferation via Wnt/β-catenin signaling.

## INTRODUCTION

The adult *Drosophila melanogaster* intestine is a powerful model to study stem cell proliferation.^1–5^ The fly gut has many important physiological functions, most notably nutrient absorption, acting as a physical barrier and providing immunity to enteric pathogens and chemical insults.^1,2^ A conserved hallmark of the gut is the dynamic nature of its architecture.^3–5^ The *Drosophila* midgut is the largest and central portion of the intestines, and it is analogous to the mammalian small intestine in function and, to some extent, cellular architecture and composition.^6^

The signature feature of the midgut is its epithelium, a single cell layer acting as a barrier to separate the lumen from internal tissues. In *Drosophila*, the midgut epithelium comprises four principal cell types. Intestinal stem cells (ISCs) can self-renew and also give rise to progenitor cells termed enteroblasts (EBs), which in turn can fully differentiate into enterocytes (ECs) that comprise the bulk of the epithelium. The specification of the fourth cell type, enteroendocrine cells (EEs), has not been fully elucidated, despite the importance of these cells in mediating important paracrine signaling events.^10–12^ EBs have been proposed to differentiate into EEs;^8,9^ however, recent studies demonstrated that ISCs, when Prospero-positive, divide into a distinct progenitor type termed pre-EEs, which subsequently differentiate into EEs.^15–17^ Disruption of ISC function can lead to either excessive proliferation or precocious differentiation, often resulting in disease.^11,12^ Therefore, a detailed understanding of the pathways and mechanisms regulating ISC proliferation is an important long-term goal with therapeutic implications.

Numerous studies have shown that Wnt/β-catenin signaling, a morphogen signaling pathway that is highly conserved in animals, promotes ISC proliferation and differentiation under both physiological conditions and upon challenges such as enteric infection or chemical insults, both of which can damage the gut epithelium. More broadly, Wnt/β-catenin signaling, also known as canonical Wnt signaling, controls diverse cellular processes during animal development and homeostasis, including stem cell maintenance, cell fate specification, neural patterning, spindle orientation, cell migration, cell polarity, and gap junction communication.^20–24^ Dysregulation of canonical Wnt signaling caused by mutations of core components of this pathway is frequently linked to birth defects and many types of cancer.^25–28^ During tissue development and homeostasis, canonical Wnt signaling is thought to be the main pathway for regulating ISC proliferation and self-renewal, which drives massive renewal processes of intestinal epithelial cells.^15–17^

Despite the fundamental importance of Wnt signaling in regulating ISC proliferation and subsequent tissue renewal in the gut epithelium,^15,16,17,18^ our understanding of how Wnt signaling in different cell types within the gut contributes to these effects on ISCs is still rudimentary. Furthermore, recent studies have elucidated roles for β-catenin-independent, or non-canonical Wnt signaling pathways, which share some upstream components in the Wnt-receiving cell but do not activate β-catenin-dependent gene expression, in regulating ISC proliferation in the *Drosophila* midgut.^19^ Thus, knowledge of which cell types exhibit canonical and non-canonical Wnt signaling that collectively contribute to ISC proliferation and thus tissue maintenance in the midgut are major fundamental and unanswered questions.

A key shared player in all Wnt signaling pathways is Dishevelled (Dsh/DVL), which is recruited to the plasma membrane upon activation of Wnt receptors and co-receptors from the Frizzled and LRP families. Such recruitment catalyzes the disassembly of a multiprotein complex that facilitates proteasomal degradation of β-catenin, enabling its accumulation and subsequent translocation to the nucleus to activate TCF/LEF-dependent gene expression in the canonical pathway.^20,21^ Dsh recruitment to the plasma membrane also activates planar cell polarity and other non-canonical Wnt pathways, including the Wnt/Ca^2+^ pathway, by activation of Frizzled receptors.^22^ Thus, regulation of Dsh levels, which occurs via the ubiquitin-proteasome system and involves the action of several distinct E3 ubiquitin ligases,^36–42^ is a key control point in all Wnt signaling pathways.

Notably, we previously discovered that a mammalian multi-subunit phosphoinositide-binding protein, pleckstrin homology domain-containing family A number 4 (PLEKHA4), promotes both Wnt/β-catenin and non-canonical Wnt signaling in human cell lines by antagonizing DVL polyubiquitination by the Cullin-3 (CUL3)–Kelch-like protein 12 (KLHL12) E3 ubiquitin ligase.^24^ We found as well that Wnt/β-catenin signaling and subsequent cell proliferation in mouse models of melanoma was dependent upon PLEKHA4 expression.^25^ In *Drosophila*, knockout of the closest fly ortholog of PLEKHA4, *kramer* (*kmr*), impairs planar cell polarity in the adult wing, larval wing imaginal disc, and pupal wing disc epithelium.^24^ The absence of any discernable defects in canonical Wnt signaling in *kmr* knockout flies led us to question whether *kmr* indeed controlled Wnt/β-catenin signaling in this organism. We proposed that the extent to which PLEKHA4/*kmr* loss affected canonical or non-canonical Wnt pathways might depend on cellular and tissue contexts, where expression of other factors downstream of DVL/Dsh might govern how tuning of DVL/Dsh levels would differentially affect outcomes from these pathways.

To test this prediction, we investigated the role of *kmr* in controlling ISC proliferation in the *Drosophila* midgut, a physiological process dependent upon canonical Wnt signaling and recently linked to non-canonical Wnt signaling pathways as well.^19^ Our experimental model involved challenge of adult flies with *E. carotovora carotovora* 15 (*Ecc*15), a gram-negative bacterium that produces non-lethal infection, to damage the midgut epithelium and induce repair pathways dependent upon ISC proliferation.^1,26^ Here, we performed global knockout and cell type-specific knockdown of *kmr* and compared effects on tissue pathophysiology to those induced by knockdown of other established components of Wnt signaling pathways. As such, our study accomplished several goals. First, we establish roles for *kmr* in controlling canonical Wnt signaling in *Drosophila*. Second, we use *kmr* as a tool to elucidate roles for canonical and non-canonical Wnt signaling within different cell types in the *Drosophila* midgut. Our studies reveal not only that *kmr*-dependent canonical Wnt signaling controls ISC proliferation in the midgut but that such signaling occurs in several cell types, including Prospero-positive cells, suggesting that EEs, which our studies support derive from pre-EE progenitors, may play an unexpectedly important role in these processes.

## RESULTS

### Expression pattern of *kramer* in the *Drosophila* midgut overlaps with zones of Wnt signaling activation

The *Drosophila* midgut comprises five major regions termed R1–R5, from anterior to posterior, based on anatomical, morphometric, and histological characterization.^27,28^ We began our study by investigating levels of Wnt signaling and *kmr* expression in these five regions (Figure S1A). Consistent with previous studies, *frizzled 3* (*fz3*), a direct target gene of canonical Wnt signaling, was expressed in gradients at the boundaries of the intestinal regions with highest expression in R1 and R2 and weak expression in R2 and R2 by examination of a Fz3-RFP transgene (Figure S1A–B).^28,29^ To examine *kramer* (*kmr*) expression, we examined the fly gut-seq dataset, which contains comprehensive RNA-seq data from each of these regions.^9^ Interestingly, *kmr* expression was highest within R1 and R5, with moderate expression in R2 and R4, a pattern that aligns with *fz3* expression (Figure S1C).^9^ Coupled with our previous finding that the human ortholog of *kmr*, PLEKHA4, promotes Wnt/ϕ3-catenin signaling,^24^ we postulated that the overlapping expression patterns of *kmr* and *fz3* in the *Drosophila* midgut supports a potential role for *kmr* in regulating Wnt signaling in the fly intestine. These findings prompted us to further investigate the potential functions of *kmr* in regulating Wnt signaling *in vivo* in the *Drosophila* midgut.

### *Kmr* is required to activate canonical Wnt signaling in the fly midgut

To investigate whether *kmr* regulates canonical Wnt signaling *in* vivo, we subjected wildtype, *kmr* knockout, and tissue-specific *kmr* knockdown fly strains to infection with a non-lethal dose of *E. carotovora carotovora 15* (Ecc15), a gram-negative bacterium that damages differentiated enterocytes, followed by recovery, which features a regenerative response.^1^ First, we examined effects on *fz3* expression the midgut in two global knockout strains generated via CRISPR/Cas9-mediated mutagenesis, *kmr*^*1*^ and *kmr*^*2*^.^24^ Because canonical Wnt signaling exhibits high activity in the R5 (posterior) region,^15^ we performed our analysis primarily in this region, unless otherwise indicated. In both unchallenged (UC) and *Ecc15*-infected groups, Fz3-RFP expression was significantly lower in both *kmr* KO strains compared to control flies (Figure 1A– D).

**Figure 1.**
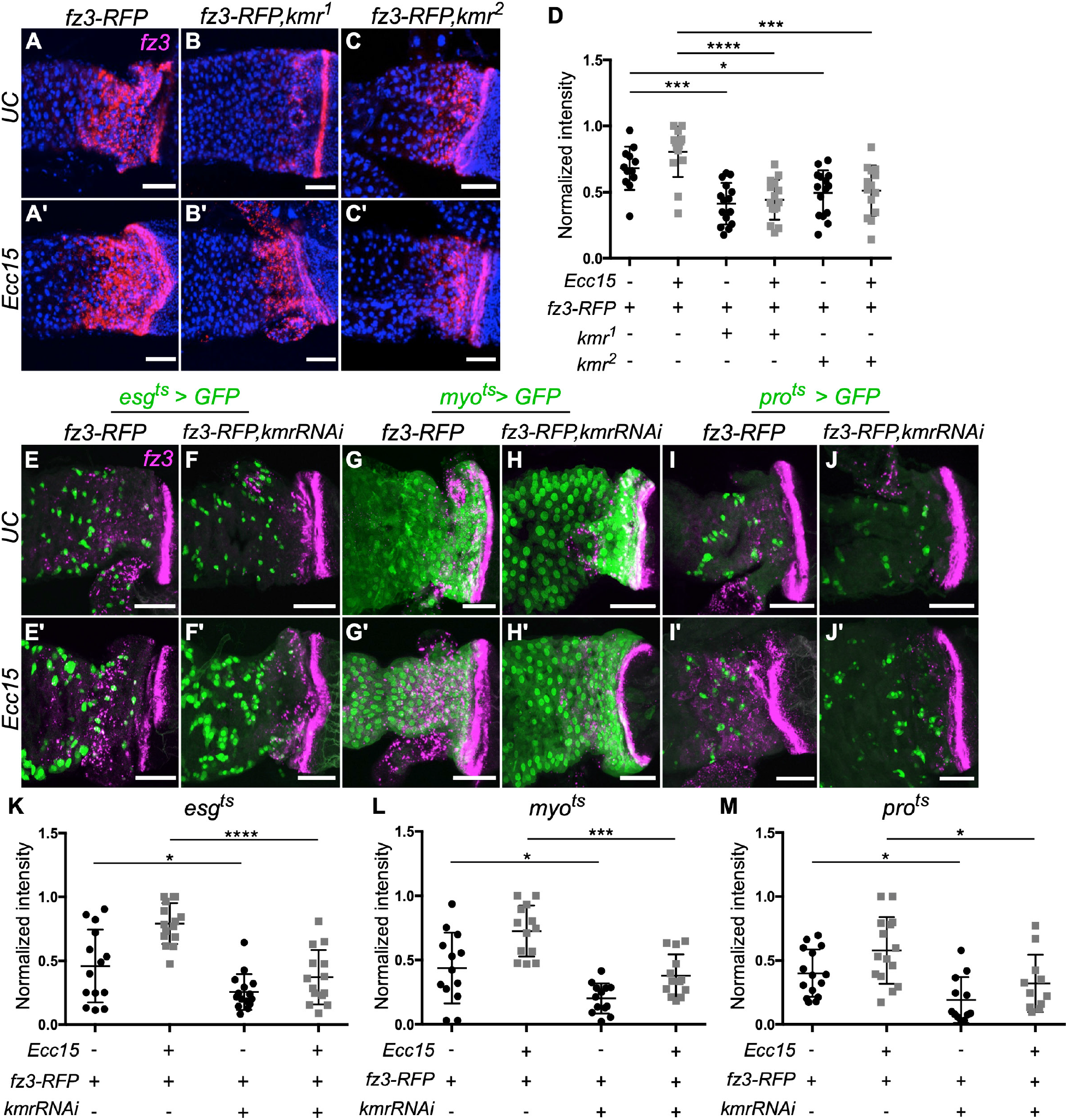
Loss of *Kramer* attenuates *frizzled3* expression in the posterior midgut. **(A–D)** Two *kmr* knockout strains (*kmr*^*1*^ and *kmr*^*2*^) exhibit decreased *fz3-RFP* expression in the posterior midgut relative to WT both in unchallenged (UC) conditions and following non-lethal infection with *Erwinia carotovora carotovora* 15 (*Ecc15*). Shown in (A–C) are representative confocal micrographs (z-projection images), with quantification for each genotype shown in (D). (E–M) Cell type-specific RNAi-mediated *kmr* knockdown decreases *fz3-RFP* expression in four different midgut cell types: intestinal stem cells (ISCs) and enteroblasts (EBs) (*esg*^*ts*^*>GFP*), enterocytes (ECs, *myo*^*ts*^*>GFP*), and enteroendocrine cells (EEs, *pro*^*ts*^*>GFP*). Shown in (E–J) are high-magnification images of the midgut-hindgut boundary, which exhibits high *fz3-RFP* expression. Green: GFP (RNAi); Magenta: Fz3-RFP (anti-mCherry antibody). Quantification of Fz3-RFP expression is shown in (K–M). Data points represent individual midguts (black circles, UC; gray squares, *Ecc15*), lines represents mean, and error bars denote standard deviation. Statistical significance was determined using one-way ANOVA with Tukey post-hoc test. ****, *p*<0.0001; ***, *p*<0.001; **, *p*<0.01; *, *p*<0.05; ns: not significant; n=11-15. Scale bars: 40 μm.

We then investigated the cellular requirements for *kmr* in governing this phenotype by performing cell type-specific *kmr* knockdown using the *Gal4/UAS-GFP* system. In ISCs and EBs, as revealed by *esg*^*ts*^*-Gal4*;*UAS-GFP* (*esg*^*ts*^*>GFP*), *kmr* knockdown downregulated the expression of *fz3-RFP* (Figure 1E–F and K). Analogously, we found that *kmr* knockdown in ECs, driven by *myo*^*ts*^*-Gal4*;*UAS-GFP* (*myo*^*ts*^*>GFP*), attenuated *fz3-RFP* expression both inside the compartments and at boundaries (Figure 1G–H and L). Finally, *kmr* knockdown in EEs, driven by *pro*^*ts*^*-Gal4*;*UAS-GFP* (*pro*^*ts*^*>GFP*), decreased *fz3-RFP* expression (Figure 1I–J and M). These studies show that *kmr* knockdown in each major midgut cell type negatively affected canonical Wnt signaling elicited by *Ecc15* challenge, consistent with its role as a positive regulator of Wnt/ϕ3-catenin signaling in mammalian cells.

### Loss of *kmr* attenuates stem cell proliferation in the midgut

We next sought to establish the role of *kmr* in governing ISC proliferation using *Ecc15* infection followed by recovery. As a primary readout, we imaged and quantified the number of dividing cells positive for phosphorylated histone 3 at serine 10 (pH3^+^).^1,9,28^ Both global *kmr* knockout strains exhibited a decreased number of pH3^+^ stem cells relative to wildtype control (Figure 2A–C and E). To further confirm the specificity of the effects to *kmr* knockout in the two global knockout strains, we generated a *kmr*^*1*^*/kmr*^*2*^ compound heterozygote and found that it displayed an identical decrease in number of pH3^+^ cells (Figure 2D–E).

**Figure 2.**
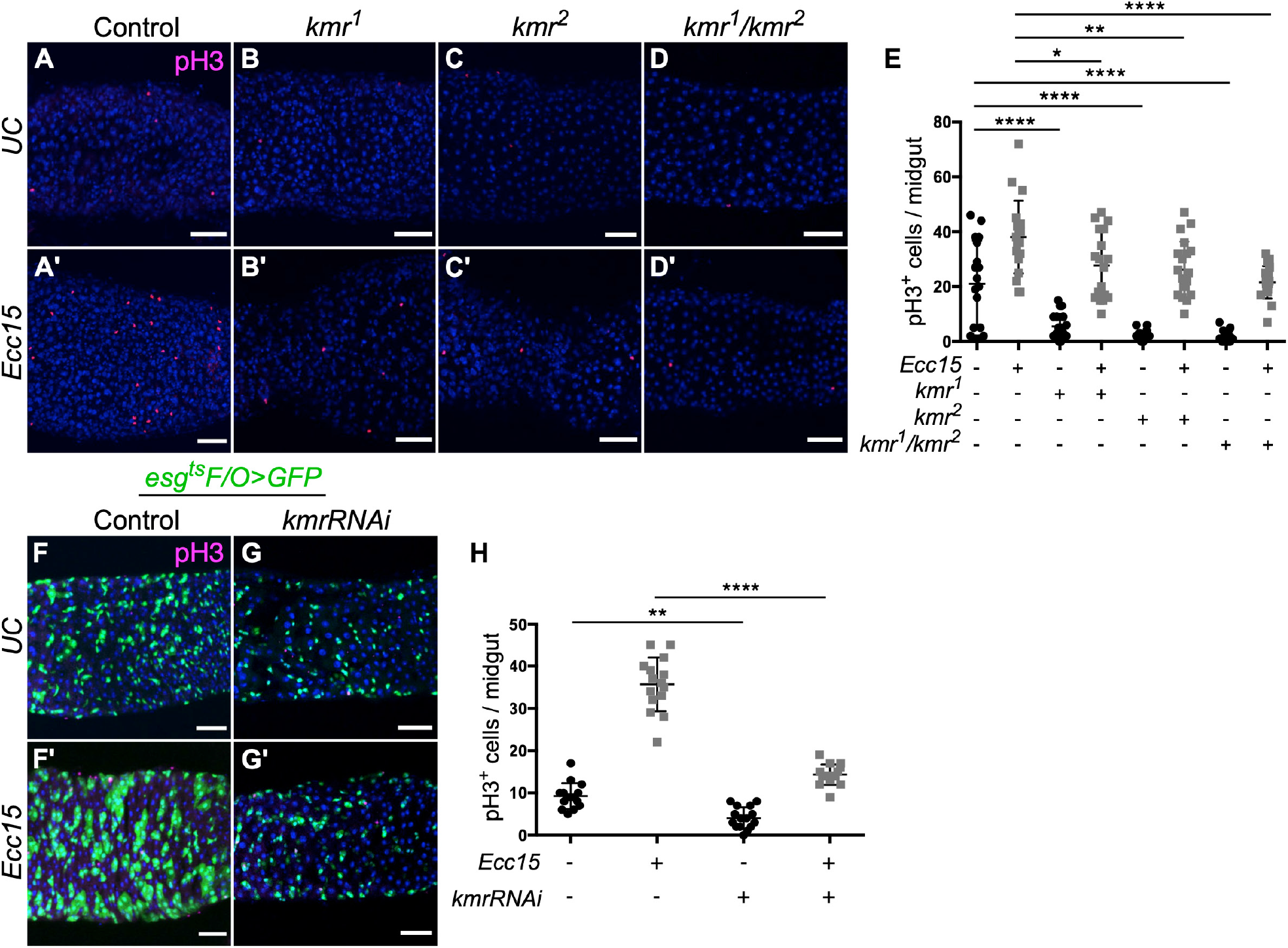
Inactivation of *kramer* downregulates intestinal stem cell proliferation. (A–E) *Kmr* knockout reduces intestinal stem cell proliferation in the midgut. Shown in (A–D) are representative confocal micrographs of number of phospho-histone H3-positive (pH3^+^) cells in the posterior midgut from unchallenged (UC) or *Ecc15*-challenged flies from the indicated genotypes: WT, *kmr* homozygous knockout (*kmr*^*1*^ and *kmr*^*2*^), and compound heterozygote (*kmr*^*1*^*/kmr*^*2*^). Quantification of number of pH3^+^ cells is shown in (E). (F–H) Conditional *kmr* knockdown in ISCs and their progeny using the *esg*^*ts*^ *F/O* system decreases number of pH3^+^ cells in unchallenged and *Ecc15*-challenged conditions. Shown in (F–G) are representative confocal micrographs of posterior midgut. Magenta: pH3; green: GFP (RNAi); blue: DAPI. Quantification of number of pH3^+^ cells from whole midguts are shown in (H). pH3^+^ cells are quantified in the whole midgut of indicated genotype. One-way ANOVA with Tukey post-hoc: ****, *p*<0.0001; **, *p*<0.01; *, *p*<0.05; ns: not significant; n=15–19. Scale bar: 40 μm.

As a more specific readout for ISC proliferation than pH3 staining, we used an inducible flip-out system under the control of *esg*^*ts*^ (esg^ts^ F/O), which results in GFP expression in ISCs, EBs, and their progeny following induction by temperature switch.^30,31,5^ Using this system, we found that RNAi-mediated *kmr* knockdown in decreased stem cell proliferation (Figure 2F–H), as assessed by quantification of both GFP^+^ and pH3^+^ cells. These results confirm that *kmr* promotes intestinal stem cell proliferation in the midgut.

### Cell type-specific *kmr* knockdown reveals role for Prospero^+^ cells in controlling ISC proliferation

To establish the cell type(s) in which *kmr* expression most strongly affects stem cell proliferation, we tested the effects of *kmr* knockdown in ISCs/EBs, ECs, and EEs separately. We found that *kmr* knockdown in ISCs and EBs decreased the number of pH3^+^ progenitor cells relative to WT controls, with significant differences observed in the *Ecc15*-challenged groups (Figures 3A–B, G and S2A–B). Knockdown of *kmr* in EC cells (Figures 3C–D, H and S2C–D) and in EE cells (Figures 3E–F, I and S2E–F) both resulted in significant decreases in pH3^+^ cells in *Ecc15*-challenged flies and also modest decreases in pH3^+^ cells in unchallenged flies. Previous studies have shown that inactivation of canonical Wnt signaling in ECs negatively impacts stem cell proliferation.^29,32,33^ However, our results here demonstrate that inactivation of canonical Wnt signaling not only in ISCs/EBs and ECs but also in EEs can also impair ISC proliferation.

**Figure 3.**
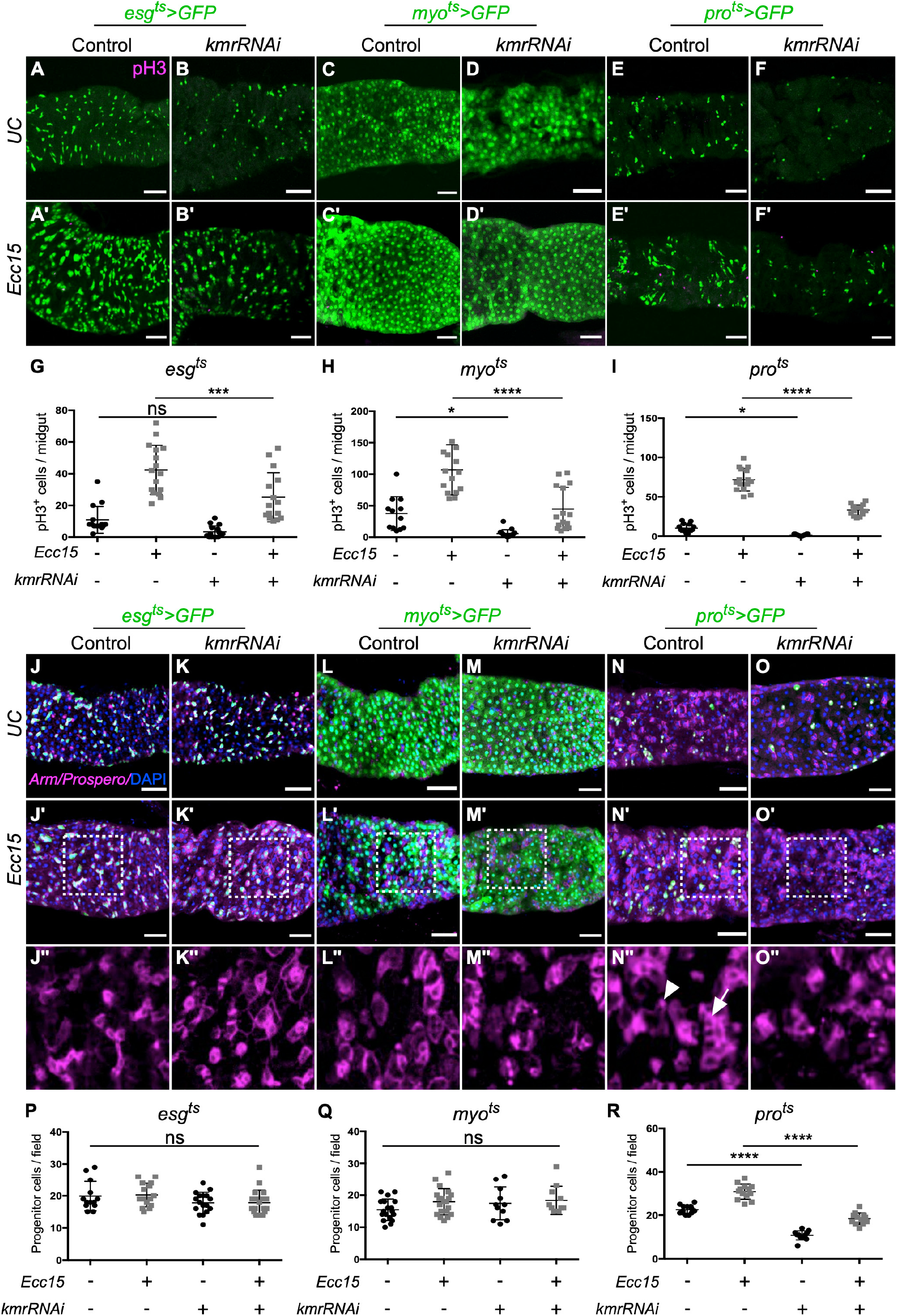
*Kramer* knockdown in multiple cell types, including enteroendocrine cells, downregulates intestinal stem cell proliferation. (A–I) RNAi-mediated knockdown of *kmr* in four different midgut cell types decreases number of pH3^+^ proliferating cells under unchallenged (UC) and *Ecc15*-challenged conditions. *Kmr* RNAi is driven by the indicated promoter and marked by GFP expression: ISCs and EBs (*esg*^*ts*^*>GFP*), ECs (*myo*^*ts*^*>GFP*), and EEs (*pro*^*ts*^*>GFP*). Shown are representative confocal micrographs of posterior midguts. Green: GFP (RNAi); magenta: pH3^+^ (proliferating cells). See also Figure S2 for monochrome images of pH3^+^ fluorescence. Quantification of number of pH3^+^ cells in the whole midgut of indicated genotype is shown in (G– I). (J–R) Knockdown of *kmr* only in EE cells, but not ISCs/EBs or ECs, causes a reduction of Armadillo^+^/Prospero^−^ (Arm^+^/Pro^−^) ISCs. Midguts from unchallenged or *Ecc15*-challenged flies of the same genotypes as above were immunostained with antibodies against Armadillo, whose cortical localization indicates progenitor cells (example shown with arrow), and Prospero, whose nuclear localization indicates EEs (example shown with arrowhead). (J’’–O”) higher maginification of the boxed areas in (J’-O’). Quantification of number of progenitor (Arm^+^/Pro^−^) cells in same boxed areas in posterior midguts is shown in (P–Q). See Figure S3 for examination of number of progenitor cells in ISCs/EBs and their progeny using the *esg*^*ts*^ *F/O* system. One-way ANOVA (Tukey post-hoc): ****, *p*<0.0001; ***, *p*<0.001; *, *p*<0.05; ns: not significant; n=11– 18. Scale bars: 40 μm.

Because of the broad nature of pH3^+^ as a marker of all proliferating cells, we next more directly tested the effects of *kmr* knockdown on stem cell proliferation using the progenitor cell marker Armadillo (Arm). Double-staining of Arm and Prospero (Arm/Pro) enables identification of both progenitor cells (ISCs and EBs), which have high levels of membrane-associated with Arm and lack of nuclear Pro staining, and pre-EE/EE cells, which exhibit strong nuclear Pro staining.^32,33^ Intriguingly, *kmr* knockdown in ISCs/EBs (Figure 3J–K and P) or in ECs (Figure 3L–M and Q) caused no change in the number of progenitor cells (Arm^+^/Pro^−^), but *kmr* knockdown in EEs (Figure 3N–O and R) resulted in significantly fewer progenitor cells compared to control, both in unchallenged and *Ecc15*-challenged flies. Consistent with these data, examination of number of progenitor cells using Arm/Pro staining following *kmr* knockdown using the esg^ts^ F/O system revealed a decrease under both unchallenged and *Ecc15*-challenged conditions (Figure S3). Together, these results suggest that *kmr* knockdown, which inactivates canonical Wnt signaling in EEs, decreases intestinal stem cell proliferation in a non-cell-autonomous manner.

### *Kmr* is required for EE differentiation and maintenance

The effects seen above of *kmr* and, by extension, Wnt signaling, in EEs came as a surprise, as there is a limited understanding of how Wnt signaling in this cell type might control ISC proliferation, prompting us to investigate this finding deeper.^8,19^ First, we hypothesized that *kmr-* dependent Wnt signaling in EEs might regulate the production and maintenance of EE cells themselves. To test this hypothesis, we examined Pro staining, as a marker of EEs, in WT and *kmr* knockout flies. We found that global *kmr* knockout decreased the number of Pro^+^ (EE) cells in both unchallenged and *Ecc15*-challenged midguts (Figure 4A–D).

**Figure 4.**
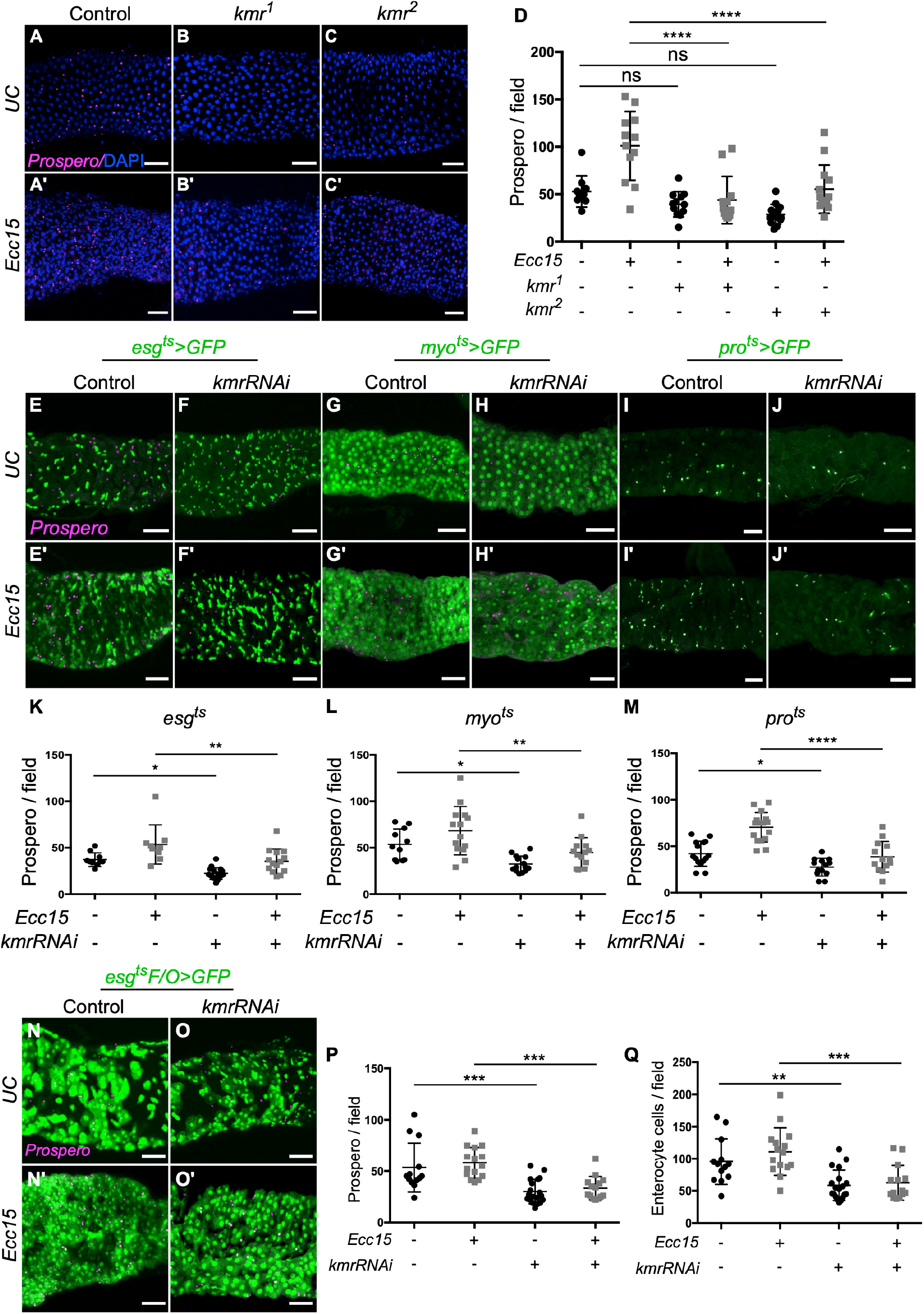
*Kramer* knockdown impairs differentiation of enteroendocrine cells. (A–D) *Kmr* knockout flies exhibit fewer EEs, marked with Prospero, compared to WT under unchallenged and *Ecc15*-challenged conditions, as shown by representative confocal micrographs in (A–C) (Magenta: anti-Prospero antibody; Blue: DAPI), with quantification of number of Pro^+^ cells shown in (D). (E–M) Cell type-specific *kmr* knockdown in four cell types in the midgut causes decrease in the number of EEs in the midgut relative to control under unchallenged and *Ecc15*-challenged conditions, as shown by representative confocal micrographs in (E–J) (Magenta: anti-Prospero antibody; Green: GFP (RNAi)), with quantification of number of Pro^+^ cells shown in (K–M). (N– Q) Conditional *kmr* knockdown in ISCs and their progeny using the *esg*^*ts*^ *F/O* system decreases number of Pro^+^ and enterocyte (EC) cells 48 h post-induction of *esg*^*ts*^ *F/O* under both unchallenged (UC) and *Ecc15*-challenged conditions. Representative images of posterior midguts are shown in N and quantification of number of Pro^+^ and ECs shown in P–Q. Pro^+^ (magenta) and ECs (green, identified by morphology) were quantified form the same fields of view. One-way ANOVA (Tukey post-hoc): ****, *p*<0.0001; **, *p*<0.01; *, *p*<0.05; ns: not significant; n=9–15. Scale bars: 40 μm.

Similar to previous experiments, we then disrupted *kmr* in different cell types in the midgut using RNAi-mediated knockdown. These studies revealed that *kmr* knockdown in ISCs/EBs (Figure 4E–F and K), ECs (Figure 4G–H and L), and EEs (Figure 4I–J and M) all reduced the number of Pro^+^ (EE) cells relative to control, in both unchallenged and *Ecc15*-challenged flies. We further used the esg^ts^ F/O system to assess the extent of ISC differentiation into EEs and ECs 48 h post-induction of *kmr* knockdown. EEs were identified using Pro staining, and ECs were identified morphologically, based on their large nuclear and cytoplasmic size, within the GFP^+^ population. We found that *kmr* knockdown reduced the number of Pro^+^ cells and ECs under both unchallenged and *Ecc15*-challenged conditions (Figure 4N–Q). These results suggest that *kmr*, and by extension *kmr*-dependent Wnt signaling, controls the differentiation and maintenance of EEs in the midgut, though an alternate interpretation is that the pool of Pro^+^ ISCs that give rise to pre-EEs are also affected by *kmr* knockdown.

### Regulation of the *kelch* E3 ubiquitin ligase adaptor by *kramer* controls proliferation in the midgut

We next set out to examine the mechanism underlying the effects of *kmr* on ISC proliferation in the midgut. We have previously established the mammalian *kmr* ortholog PLEKHA4 as a positive regulator of DVL levels via sequestration and inactivation of the Cullin-3 substrate adaptor Kelch-like protein 12 (KLHL12) within clusters, preventing CUL3–KLHL12-mediated polyubiquitination of DVL, a key activatior of Wnt/β-catenin and non-canonical Wnt signaling ^24^. Therefore, we reasoned that the effects of *kmr* on ISC proliferation in the midgut might involve antagonism of a fly ortholog of KLHL12. We therefore investigated the genetic interaction of *kmr* with each of the two closest KLHL12 orthologs, *kelch (kel)* and *diablo (dbo)*. We first assessed ISC proliferation by quantifying the number of pH3^+^ cells in WT flies compared to those expressing cell type-specific *kel* RNAi in either ISCs/EBs (Figures 5A–E and S4A–D), ECs (Figures 5F–J and S4E–H), or EEs (Figures 5K–O and S4I–L).

**Figure 5.**
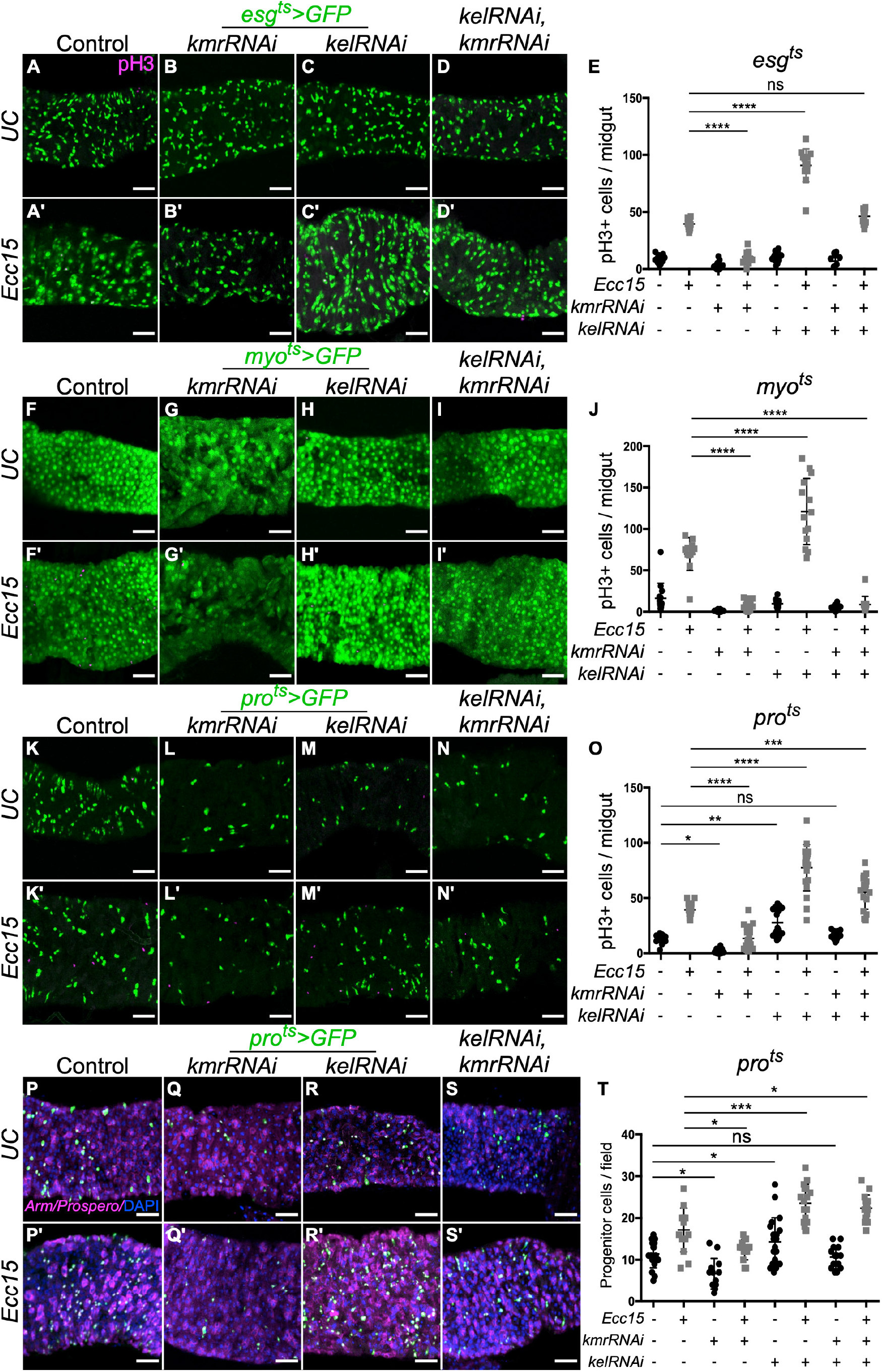
Knockdown of *kelch* partially restores the intestinal stem cell proliferation defects caused by *kramer* downregulation in midgut cells. (A–O) Examination of number of pH3^+^ cells in the midgut in unchallenged or *Ecc15*-challenged flies expressing conditional, cell type-specific knockdown of *kmr* and/or the Cullin-3 E3 ligase substrate-specific adaptor *kelch* (*kel*) in ISCs/EBs (A–E, *esg*^*ts*^*>GFP*), ECs (F–J, *myo*^*ts*^*>GFP*), or EEs (K–O, *pro*^*ts*^*>GFP*). Shown for each set are representative confocal micrographs (magenta: pH3; green: GFP (RNAi)) and quantification of number of pH3^+^ cells in the whole midgut. See also Figure S4 for monochrome images of pH3 staining and Figure S5 for examination of effects of the *kel* paralog *diablo* (*dbo*). (P–T) Examination of number of Arm^+^/Pro^−^ progenitor cells in the midgut in unchallenged or *Ecc15*-challenged flies expressing conditional, cell type-specific knockdown of *kmr* and/or *kel* in EEs. Shown are representative confocal micrographs (magenta: Arm/Pro; green: GFP (RNAi)) and quantification of number of Arm^+^/Pro^−^ cells in the same field of posterior midguts. See Figure S6 for similar analysis of Arm^+^/Pro^−^ cells in ISC/EB or EC-specific knockdown of *kmr* and/or *kel*. One-way ANOVA (Tukey post-hoc): ****, *p*<0.0001; ***, *p*<0.001; **, *p*<0.01; *, *p*<0.05; ns, not significant; n=12–18. Scale bars: 40 μm.

These studies revealed several findings. First, by comparing *kel* knockdown to control in *Ecc15*-challenged flies, it is apparent that *kel* knockdown in all cell types significantly increased total cell proliferation in the midgut, consistent for a role of *kel* as a suppressor of Wnt/β-catenin signaling via its polyubiquitination of DVL (ISCs/EBs: Figure 5C and E; ECs: Figure 5H and J; EEs: Figure 5M and O).^24^ The effect of *kel* RNAi in EE cells on this phenotype was strong enough to elicit a statistically significant increase in proliferating cells even in unchallenged flies (Figure 5M and O).

Second, we performed comparison of *kmr/kel* double knockdown flies to the three other groups (control and single knockdown of either *kmr* or *kel*). When such knockdown was performed in ISCs/EBs (Figure 5A–E) and in EEs (Figure 5K–O), the *kmr/kel* double knockdown midguts exhibited an intermediate extent of proliferation: lower than *kel* knockdown, higher than *kmr* knockdown, and comparable to control. One interpretation of these results is that the addition of *kel* knockdown in the double knockdown strain significantly diminished the effects of single *kmr* knockdown on proliferation, consistent with a role for *kmr* as an inhibitor of *kel* as demonstrated biochemically for their mammalian orthologs, PLEKHA4 and KLHL12.^24^ Furthermore, we found no effects of knockdown of *dbo*, the other potential KLHL12 fly ortholog, on this phenotype and no evidence for genetic interaction with *kmr* (Figure S5), further supporting a relationship between *kmr* and *kel* in controlling Wnt signaling in this tissue, consistent with our earlier studies.

However, the observed effects on proliferation were partial, i.e., proliferation in the double knockdown strain was not as high as *kel* knockdown alone. Though incomplete knockdown is a possible explanation for this intermediate result, the data are consistent with several types of genetic relationships between *kmr* and *kel*, critically they are identical to those that we observed in mammalian cells using RNAi-mediated knockdown of the orthologs of these genes, PLEKHA4 and KLHL12, alone or in combination.^24^ Therefore, these results strongly suggest that the mechanism connecting *kmr* and *kel* in these cells is similar to that established by us and others as regulators of Wnt signaling pathways via effects on DVL/Dsh ubiquitination.^24,34,35^ In ECs, we found that *kmr/kel* double knockdown nearly completely eliminated proliferation following *Ecc15* challenge, similar to *kmr* single knockdown, suggesting a different relationship between these genes in this cell type and one that may warrant further study (Figure 5F–J).

As a more precise measure of effects of *kmr* and/or *kel* knockdown on ISC proliferation, we performed Arm/Pro staining on the double knockdown strains, as before (Figure 3). Our earlier results showed that *kmr* knockdown exclusively in EEs attenuated ISC proliferation (Figure 3N–O and R). Here, we asked whether *kel* expression in EEs also regulates ISC proliferation, and whether *kel* knockdown could restore the reduction of ISC proliferation induced *kmr* RNAi. As expected, given the lack of an effect of *kmr* knockdown in ISCs/EBs and ECs on this phenotype (Figure 3), individual knockdown of *kel* or combined *kmr/kel* double knockdown also had no effect on the number of Arm^+^/Pro^−^ progenitors (Figure S6). By contrast, *kel* knockdown in EEs induced overproliferation of Arm^+^/Pro^−^ progenitors (Figure 5P–T) compared to control and fully eliminated the effect of *kmr* knockdown in reducing ISC proliferation. Taken together, these results indicate that *kmr* and *kelch* regulate ISC proliferation in the midgut in a manner consistent to previous mechanistic findings in mammalian cells and other tissues in adult and larval *Drosophila*, where PLEKHA4/*kmr* was found to enhance both canonical Wnt/β-catenin and PCP, a non-canonical Wnt signaling pathway.^24^

### *Kmr* controls ISC proliferation via effects on both canonical and non-canonical Wnt signaling

Because PLEKHA4/*kmr* regulates DVL/Dsh levels, which can affect both canonical and non-canonical Wnt signaling pathways, a key outstanding question is whether the effects of *kmr* observed here on ISC proliferation in the midgut are due entirely to Wnt/β-catenin signaling, PCP or another non-canonical pathway, or a combination. Resolving this distinction is important because, though previous work from our lab established PLEKHA4 as a regulator of Wnt/β-catenin signaling in mammalian cells, in *Drosophila*, we found it to regulate hair polarization, a hallmark PCP phenotype, in several tissues, consistent with previous studies implicating *kel* and *dbo* in PCP signaling.^35^ To date, no studies have found that *kmr* controls Wnt/β-catenin signaling.

*Kmr* knockdown globally and in relevant midgut cell types individually all caused a decrease in expression of the *fz3-RFP* reporter of canonical Wnt signaling, indicating that *kmr* is indeed capable of regulating this pathway in the midgut (Figure 1). However, whether the observed effects of *kmr* on ISC proliferation elsewhere in this study (Figures 2–5) are due to effects on canonical Wnt signaling and/or PCP remain undetermined. To distinguish between these possibilities, we inactivated each of those pathways individually in ISCs/EBs, ECs, or EEs, to determine which perturbation would phenocopy *kmr* knockdown. To target PCP, we used knockdown of *otk*, the *Drosophila* ortholog of PTK7, a cell-surface receptor that regulates PCP,^19^ and to inactivate canonical Wnt/β-catenin, we performed knockdown of *pangolin* (*pan*), the *Drosophila* TCF/Lef ortholog that acts as a transcription factor downstream of Arm (β-catenin).^36^

We first evaluated overall proliferation in the midgut using pH3 staining (Figures 6A–L and S7). We found that in *Ecc15*-challenged flies, *otk* knockdown in ISCs/EBs (Figure 6A–B and D) and in EEs (Figure 6I–J and L) but not in ECs (Figure 6E–F and H) reduced overall proliferation. By contrast, *pan* knockdown in all cell types significantly reduced proliferation (ISCs/EBs: Figure 6A and C–D; ECs: Figure 6E and G–H; EEs: Figure 6I and K–L). These results indicate that *pan* knockdown exactly phenocopies *kmr* knockdown in regulation of stem cell proliferation, suggesting that *kmr* acts on these phenotypes through canonical Wnt signaling. At the same time, these data indicate important roles for Wnt/β-catenin sigaling not only within ISCs/EBs and ECs, as was previously known, but also in EEs, in controlling proliferation. The effects of *otk* knockdown on proliferation partially phenocopies *kmr* knockdown, suggesting that *kmr* regulation of PCP may be relevant within progenitors, consistent with its role in regulation of PCP in other tissues.^24^ The *otk* knockdown experiments confirm recent studies showing a role for PCP within ISCs for controlling proliferation,^19^ and they also reveal unknown potential roles for PCP within Pro^+^ cells in regulating proliferation.

**Figure 6.**
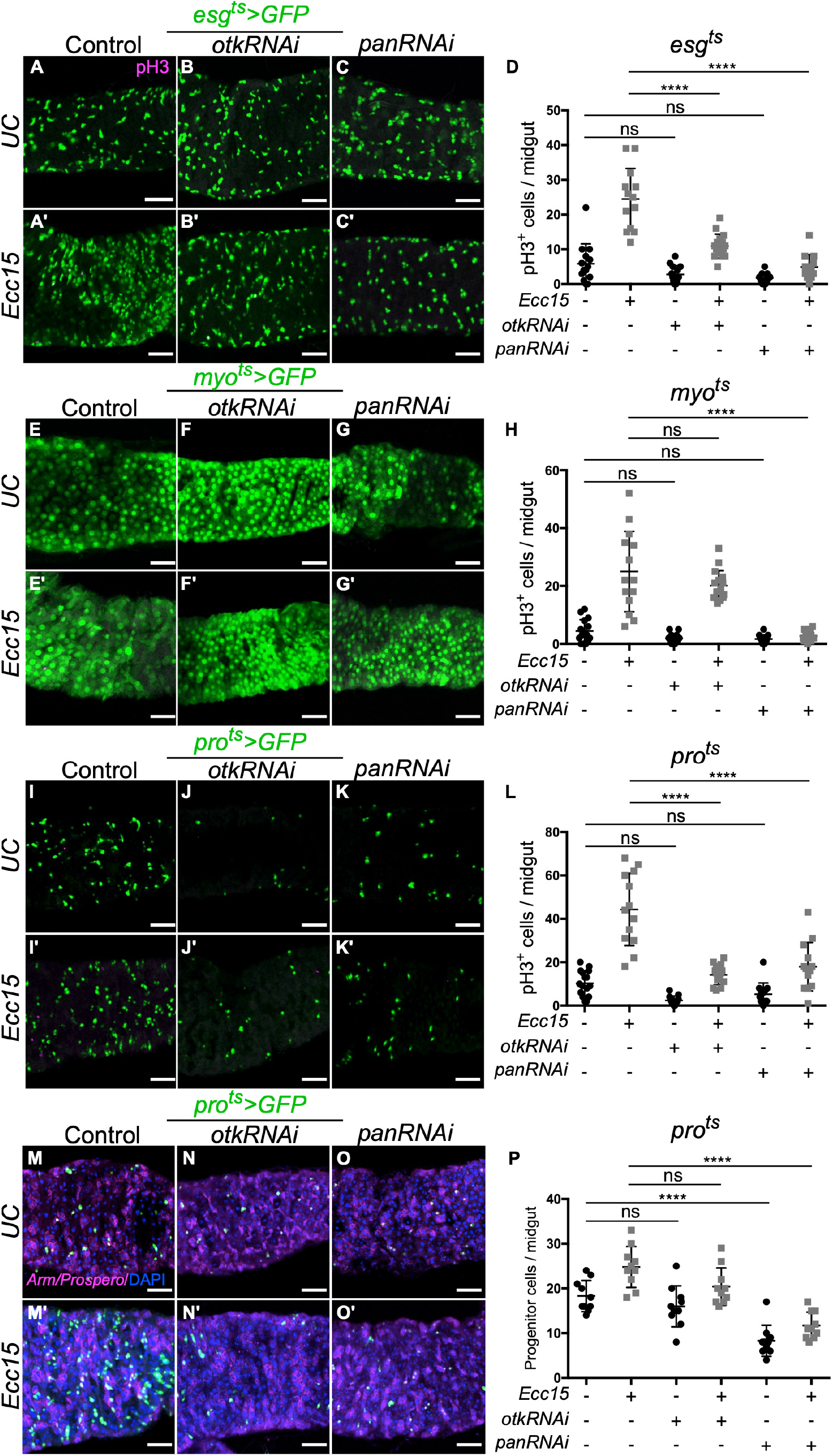
Suppression of Wnt/β-catenin signaling using *pangolin* (*pan*) knockdown, but not of planar cell polarity using *off-track* (*otk*) knockdown, phenocopies effects of *kmr* knockdown on intestinal stem cell proliferation in the midgut. (A–L) Examination of number of pH3^+^ cells in the midgut in unchallenged or *Ecc15*-challenged flies expressing conditional, cell type-specific knockdown of *pan* or *otk* in ISCs/EBs (A–D, *esg*^*ts*^*>GFP*), ECs (E–H, *myo*^*ts*^*>GFP*), or EEs (I–L, *pro*^*ts*^*>GFP*). Shown for each set are representative confocal micrographs (magenta: pH3; green: GFP (RNAi)) and quantification of number of pH3^+^ cells. See also Figure S6 for monochrome images of pH3 staining. (M–P) Examination of number of Arm^+^/Pro^−^ progenitor cells in the midgut in unchallenged or *Ecc15*-challenged flies expressing conditional, cell type-specific knockdown of *pan* or *otk* in EEs. Shown are representative confocal micrographs (magenta: Arm/Pro; green: GFP (RNAi)) and quantification of number of Arm^+^/Pro^−^ cells. One-way ANOVA (Tukey post-hoc): ****, *p*<0.0001; ns, not significant; n=11–14. Scale bars: 40 μm.

Finally, we used Arm/Pro staining to directly quantify effects of *otk* or *pan* knockdown in EEs on ISC/EB proliferation (Figure 6M–P), given the strong effects of *kmr* knockdown in these cells on this phenotype (Figure 3N–O and R). We found that *otk* knockdown in EE cells had no effect on the number of Arm^+^/Pro^−^cells both in unchallenged conditions and following *Ecc15* challenge (Figures 6M–N and P). By contrast, *pan* knockdown impaired stem cell proliferation by this readout compared to control, both in unchallenged and *Ecc15*-challenged flies (Figures 6M and O–P). These results strongly suggest that *kmr* within Pro^+^ cells regulates stem cell proliferation in the midgut via canonical Wnt signaling. More broadly, our results implicate *kmr* as a new regulator of canonical Wnt signaling in the fly midgut via effects in several pathways, and our data suggest unexpected roles for Wnt signaling within EEs in the regulation of stem cell proliferation in the construction of the gut epithelium in *Drosophila* (Figure 7).

**Figure 7.**
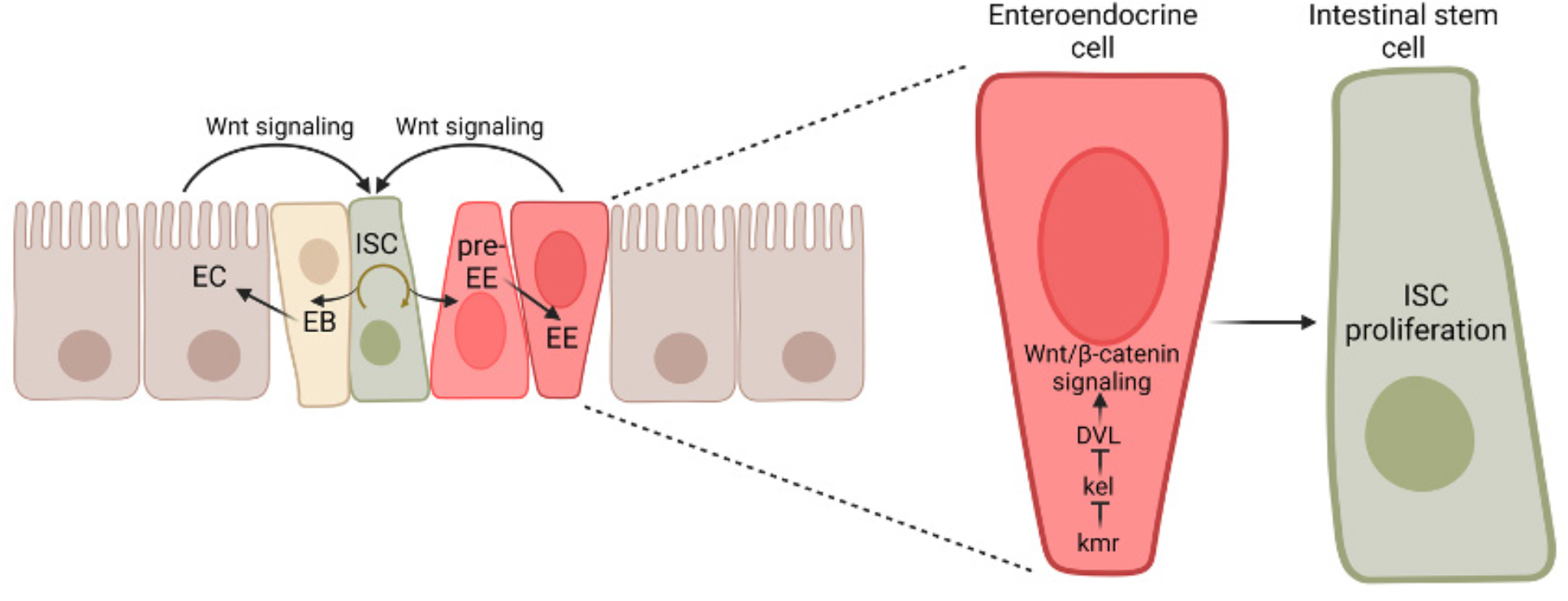
Working model for *kramer* function in the *Drosophila* midgut. Intestinal epithelial homeostasis requires continuous ISC self-renewal and differentiation into other cell types, including enteroblasts (EB) that give rise to enterocytes (EC), and enteroendocrine cells (EEs) that derive from a distinct progenitor termed pre-EEs. In turn, these multiple cell types can provide signaling including secreted Wnt ligands to sustain ISC proliferation. In this model, canonical Wnt/β-catenin signaling within multiple cell types, including enteroendocrine cells, promotes stem cell proliferation in the midgut. Such signaling is dependent upon *kramer*, which acts by antagonizing *kelch*, a negative regulator of DVL/Dsh and Wnt signaling pathways.

## DISCUSSION

Proliferation and differentiation of intestinal stem cells in the adult *Drosophila* midgut is essential to maintain the balance of tissue homeostasis and prevent excessive proliferation in this tissue. Wnt/β-catenin signaling plays central roles in tissue maintenance during development, including in the midgut. However, how canonical Wnt signaling is activated within intestinal stem cell progenitors and by fully differentiated progeny cells is still not well understood. Here, we discovered a new player, *kramer (kmr)*, that regulates canonical Wnt signaling in the *Drosophila* midgut using a challenge with the non-lethal pathogen *Erwinia carotovora carotovora* (*Ecc15*) to induce massive ISC proliferation required to rebuild the damaged gut epithelium.

Using this system, we established *kmr* as a positive regulator of ISC proliferation and Wnt/β-catenin signaling. We then used inducible, cell type-specific *kmr* knockdown as a tool to interrogate the requirements for Wnt signaling within each cell type in the gut for controlling ISC proliferation. In addition to known effects of Wnt signaling within intestinal stem cells (ISCs), their immediate downstream progenitors termed enteroblasts (EBs), and fully differentiated enterocytes (ECs) in governing this process, our study unexpectedly points to roles for Wnt signaling within enteroendocrine cells (EEs) as a mechanism controlling stem cell proliferation in the *Drosophila* midgut.

We first showed that interruption of *kmr* function decreases the expression of a canonical Wnt signaling target gene, *fz3*, in the posterior midgut, consistent with studies demonstrating *fz3RFP* expression in both ISCs and ECs showing that knockout of Wnt signaling components led to loss of *fz3* expression in midgut region R5.^29^ Though ECs are the primary cell type in which Wnt signaling is activated,^28,29,32^ our studies using *kmr* knockdown in multiple cell types suggest that Wnt signaling is also active in ISCs/EBs and EEs in posterior end of the midgut. These studies also revealed that loss of *kmr* in multiple cell types results in fewer EE cells and decreased ISC proliferation in the posterior midgut, indicating a role for *kmr* in EE differentiation.

Further, *kmr* knockdown in all cell types caused a decrease in proliferating, phospho-H3 positive (pH3^+^) cells, though staining with Armadillo/Prospero antibodies, which enables identification of ISCs/EBs and EEs, suggests that *kmr* knockdown in EE cells specifically downregulates stem cell proliferation. This distinction is important because pH3^+^ cells include all dividing progenitor cells, including ISCs/EBs and pre-EEs, and our data showed that *kmr* knockdown in all cell types significantly decreased dividing stem cells. Notably, *kmr* knockdown in ISCs/EBs and in ECs did not result in a defect in number of Arm^+^/Pro^−^ cells (i.e., ISCs/EBs); However, *kmr* knockdown in Pro^+^ cells led to a reduction in number of ISCs/EBs, suggesting that *kmr* expression in EE cells regulates the proliferation of neighboring ISCs in a non-autonomous manner. These data also provide support to a model wherein EEs derive not from EBs^8,9^ but instead from a distinct set of progenitors termed pre-EEs.^10,37,38^

Our study also sheds light on the mechanisms by which *kmr* expression in distinct midgut cell types might regulate the physiological response to *Ecc15* infection. Prior work on the mammalian ortholog of *kmr*, PLEKHA4, revealed that it promotes Wnt signaling pathways via physical interaction with KLHL12, a CUL3 E3 ubiquitin ligase substrate-specific adaptor and negative regulator of DVL.^24^ By binding to KLHL12 and sequestering it in plasma membrane-associated clusters, PLEKHA4 prevents DVL polyubiquitination by CUL3–KLHL12, ultimately causing elevated DVL levels and enhanced Wnt signaling in Wnt-receiving cells.

Because of the pleiotropic roles for DVL in both canonical Wnt/β-catenin and non-canonical β-catenin-independent pathways,^22^ we found that PLEKHA4 knockdown affected both of these pathways.^24^ Our previous studies on *kmr* in *Drosophila*, focusing on hair patterning, identified defects in planar cell polarity (PCP), a Frizzled- and Dsh-dependent pathway in flies.^24^ However, the present study identifies for the first time a role for *kmr* in promoting canonical Wnt/β-catenin signaling in vivo. By analyzing knockdown of the two fly KLHL12 orthologs, *kelch* and *diablo*, we found that *kmr* acts in opposition to *kelch*, much as PLEKHA4 does with KLHL12 in mammalian cells, supporting the established mechanism of action. Finally, we independently blocked Wnt/β-catenin and PCP signaling and assessed ISC proliferation following *Ecc15* challenge and found that only blockade of Wnt/β-catenin signaling exactly phenocopied loss of *kmr*.

In conclusion, our study reveals that the Dsh regulator *kmr* is a new, important regulator of Wnt/β-catenin signaling and ISC proliferation in vivo. As well, we harness *kmr* as a tool to systematically investigate the role of Wnt signaling in each of the major cell types in the *Drosophila* midgut in controlling ISC proliferation and differentiation during epithelial repair following challenge with an enteric pathogen. These studies suggest that Wnt signaling within enteroendocrine cells can control this process, adding a new layer of regulation to this important physiological process. Future studies will be necessary to identify downstream mechanisms by which EEs may non-autonomously regulate ISC proliferation in the midgut, as well as the existence of similar pathways in mammalian systems.

## MATERIALS AND METHODS

### Fly stock generation and maintenance

Flies were maintained at room temperature in standard yeast glucose medium (50 g/L yeast, 60 g/L yellow cornmeal, 40 g/L sucrose, 7 g/L fly agar, 26.5 mL/L moldex, 12 mL/L acid mix). Fly stocks used in this study include: *w*^*1118*^ and CantonS; *Gal4* drivers used were ‘*w*^*-*^; *esg-Gal4; UAS-GFP, tub-Gal80*^*ts*^*’*(*esg*^*ts*^, progenitor-specific);^39^ ‘w^*-*^; *myo1A-Gal4; UAS-GFP, tub-Gal80*^*ts*^*’* (*myo*^*ts*^, EC-specific);^4^ ‘w^*-*^; *pro-Gal4; UAS-GFP, tub-Gal80*^*ts*^*’* (*pro*^*ts*^, EE-specific);^9^ ‘*esg-Gal4; UAS-GFP, tub-Gal80*^*ts*^; *Act>STOP>Gal4,UAS-flp’* (*Esg*^*ts*^ *F/O*, progenitors^+^ marked lineage);^40^ *UAS*-*kmrRNAi* (BDSC, 51917); *UAS*-*kelchRNAi* (BDSC, 55612); *UAS*-*panRNAi* (BDSC, 26743); *UAS*-*OTKRNAi* (BDSC, 25790); *fz3-RFP* (obtained from Yashi Ahmed).^29^ Knockout strains *kmr*^*1*^ and *kmr*^*2*^ were generated with CRISPR/Cas9 gene deletion by our lab.^24^ Strains *sp/cyow; TM2/TM6B* (obtained from Chun Han) were used to cross with *kmr*^*1*^, *kmr*^*2*^, *kmrRNAi* and/or *kelchRNAi* separately, then back crossed with cell type specific Gal4 system. See Table S1 for details.

### Infection of flies with *Ecc15*

Flies were maintained at room temperature or at 18 ºC in a 12 h light/12 h dark cycle incubator. Only posterior midguts of female flies were analyed in this study, except that pH3^+^ cells were quantified from the whole midguts. CantonS, a wild type inbred line, was used as the wild type control for all experiments, except for experiments using *kramer* mutants, which used *w*^*1118*^ as control. F1 developing progenies of crosses involing the temperature-sensitive (*Gal4-Gal80*^*ts*^) system were maintained at 18 ºC, and adults were collected on the 5th day after eclosion, to allow for proper midgut development. F1 progenies were then transferred to 29 ºC for 6 days to allow for transgene induction to take effect. F1 flies were then transferred to empty vials for a 2 h starvation, followed by 12-16 h *Ecc15* infection at 29 ºC, as previously described.^1^ *Ecc*15 was cultured in 500 mL of LB medium for 16 h at 30 ºC. The culture was pelleted, resuspended in water to an OD_600_ of 200. Then *Ecc15* was diluted with the same volume of sucrose (1:4 ratio of 25% sucrose:water) to OD_100_ in a 2.5% sucrose solution. Then, 150 µL of diluted *Ecc15* culture was deposited on a Whatman filter paper, which was inserted into the food vial and placed on top of the food. In the *Ecc15* infection experiments, 2.5% sucrose was used as unchallenged control. Whole guts were dissected from *Ecc15*-infected and control flies for further analyses.

### Immunohistochemistry and histology

The immunohistochemistry protocol was adapted from a previous study ^5^. Immunostaining was performed at room temperature. *Drosophila* midguts were dissected in 1X PBS. Dissected midguts were fixed in 4% paraformaldehyde (Electron Microscopy Sciences, Hatfield, PA, Cat# 15714) for 60–90 min and subsequentially washed three times with 0.1% Triton X-100 (Cat# IB07100, IBI Scientific, Dubuque, IA) that was diluted in 1X PBS. Fixed midguts were blocked for 1 h in blocking solution (1% bovine serum albumin, Cat# A8806, Sigma-Aldrich, St. Louis, MO; 1% normal donkey serum, Cat# 102644-006, VWR, Radnor, PA. in PBS). Then the primary antibody (1:1000) was added into the blocking solution with midguts and incubated overnight. The primary antibodies used were mouse anti-pH3 (1:500, Cat# 9706S, Cell Signaling Technology, Danvers, MA), mouse anti-Armadillo (1:10, Cat# N2 7A1, DSHB), and mouse anti-Prospero (1:100, Cat# MR1A, DSHB). After three washes with 0.1% Triton X-100 in PBS, the midguts were incubated with secondary antibody (donkey anti-mouse Alexa 594 (1:400, Cat# A21203, Thermo Fisher)) for 2 h in blocking solution. Midguts were then washed three times in 1X PBS. DNA was then stained with DAPI and midguts were mounted with ProLong Diamond Antifade Mountant (Cat# P36971, Thermo Fisher) overnight. Imaging was performed on a Zeiss LSM 800 confocal laser scanning microscope equipped with EC Plan-Neofluar 10X 0.3 NA and Plan-Apochromat 20X 0.8 NA air objectives, Plan Apochromat 40X 1.4 NA and Plan-Apochromat 63X 1.4 NA f/ELYRA oil immersion objectives, 405, 488, 561, and 640 nm solid-state lasers, two GaAsP PMT detectors, and an Airyscan module (Carl Zeiss Microscopy, Thornwood, NY). Images were acquired using Zeiss Zen Blue 2.3 and analyzed using ImageJ/FIJI.^41^ pH3^+^ cells were counted directly in epifluorescence mode using a 20X objective lens. Armadillo^+^/Prospero^−^ labeled progenitor cells and Prospero^+^ cells were counted manually within a field of vision of posterior midguts.

### Statistical analysis

All plots were generated using GraphPad Prism v.7.0 (https://www.graphpad.com). Statistical significance between different treatments were assessed using one-way ANOVA followed by a Tukey’s post-hoc test.

## ACKNOWLEDGMENTS

We acknowledge support from the NIH (J.M.B.: R01GM131101; N.S.B.: R01AI148529, R21AG065733, and R01AI148541), the NSF (N.S.B.: IOS1656118), and the Sloan Foundation (J.M.B.: Sloan Research Fellowship). We thank Yashi Ahmed (Dartmouth) for providing *fz3-RFP* flies, Lin Luan for technical assistance, the Han lab for equipment, and Chun Han and members of the Baskin lab for helpful discussions.

## AUTHOR CONTRIBUTIONS

Conceptualization: H.S., A.S., A.B., N.S.B., J.M.B.; Funding Acquisition: J.M.B., N.S.B.; Investigation: H.S., A.S., A.B.; Project Administration: J.M.B.; Supervision: J.M.B.; Writing – original draft: H.S., J.M.B.; Writing – review & editing: H.S., A.S., A.B., N.S.B., J.M.B.

## SUPPLEMENTAL FIGURES

**Figure S1.**
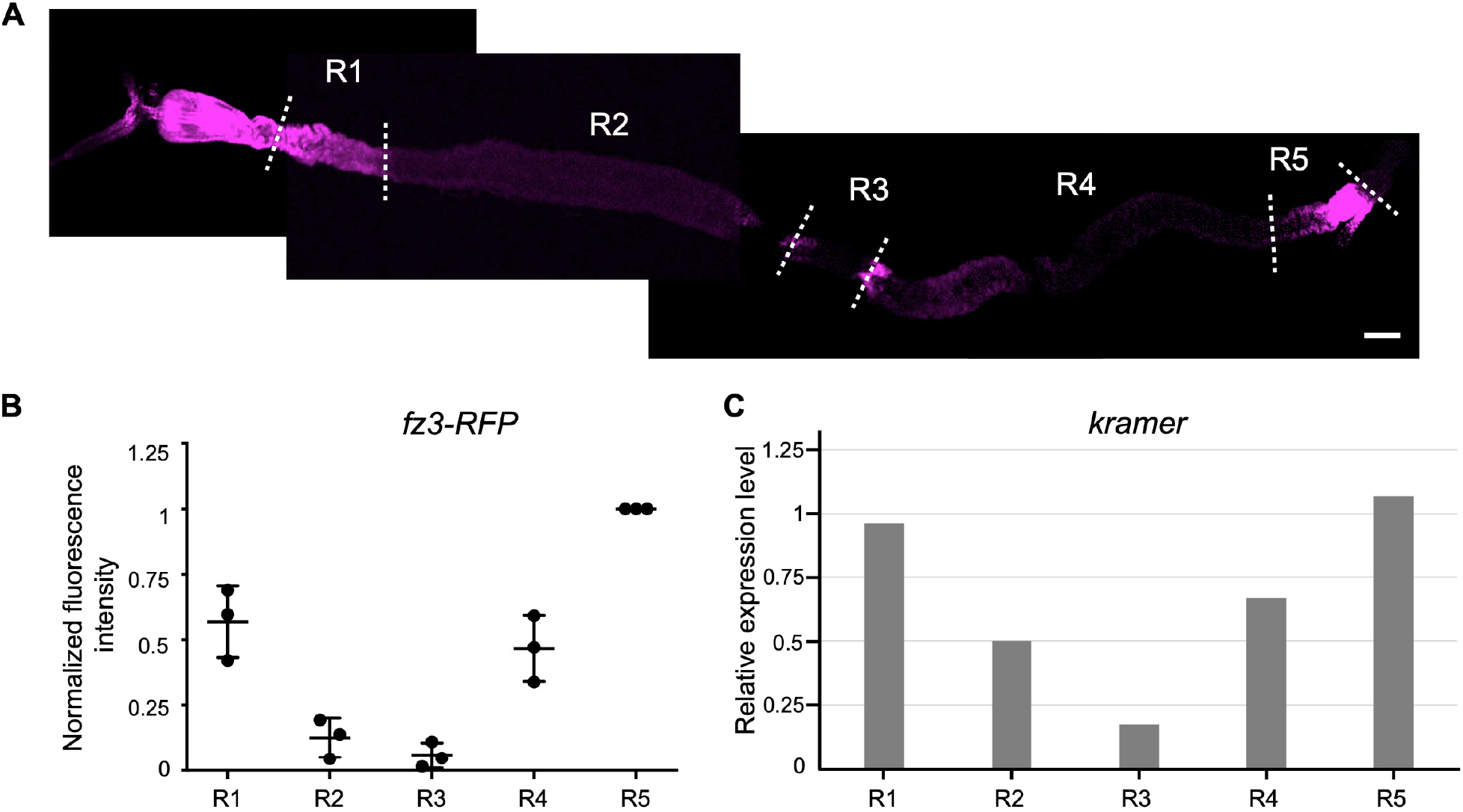
*Kramer* expression pattern in the adult *Drosophila* midgut mirrors that of the Wnt reporter *frizzled3*. (A) Immunofluorescence staining of the midgut from adult *frizzled3 (fz3)-RFP* transgenic *Drosophila*, with five regions (R1–5) indicated. Magenta: *fz3*-*RFP*, stained with mCherry antibody. Scale bar: 200 μm. (B) Quantification of *fz3*-*RFP* staining from (A). Fluorescence intensities are normalized to R5 intensity. (C) Expression levels of *kramer* (*kmr*) mRNA in different regions of adult *Drosophila* midgut. Data were obtained from FlyGut-seq as normalized expression rpkm (reads per kilobase million).

**Figure S2 (related to Figure 3).**
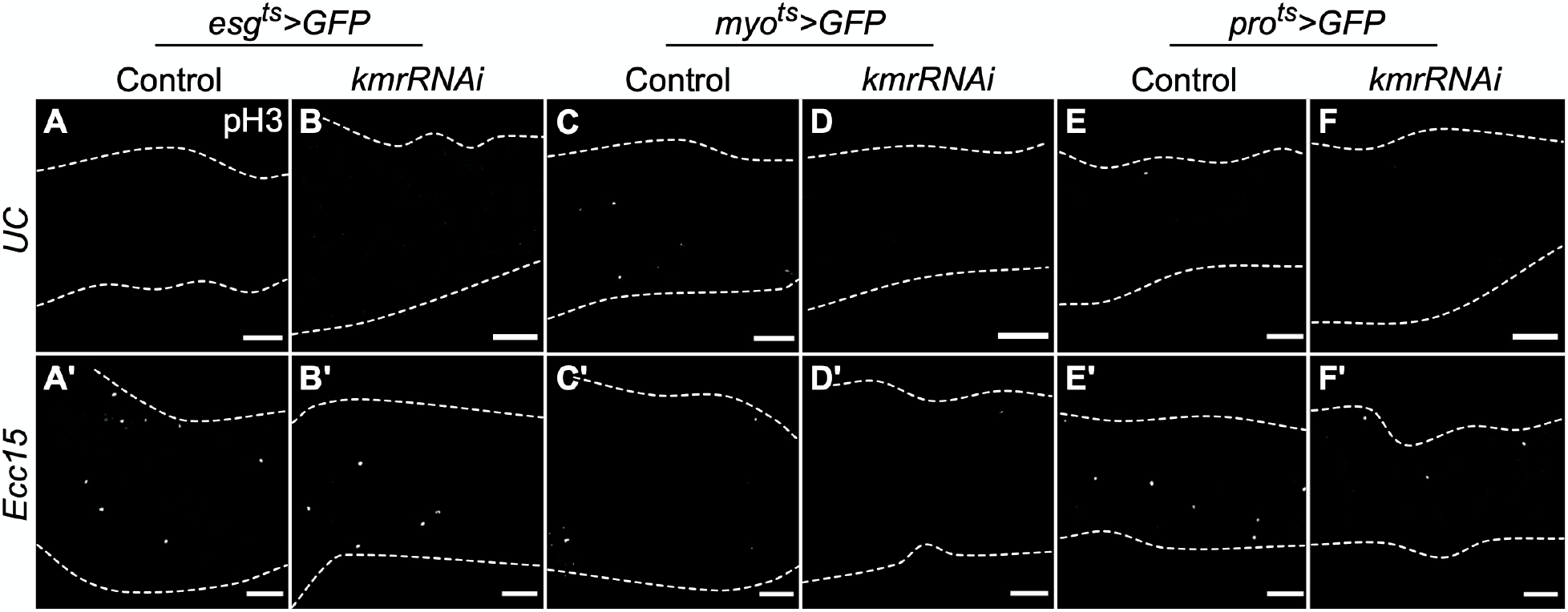
Monochrome images of pH3 staining for studies showing *kramer* knockdown in multiple cell types, including enteroendocrine cells, downregulates intestinal stem cell proliferation. Shown are monochrome images of pH3 fluorescence in cells stained as described in Figure 3A–F. Dashed lines indicate the boundary of the gut. Scale bars: 40 μm.

**Figure S3 (related to Figure 3).**
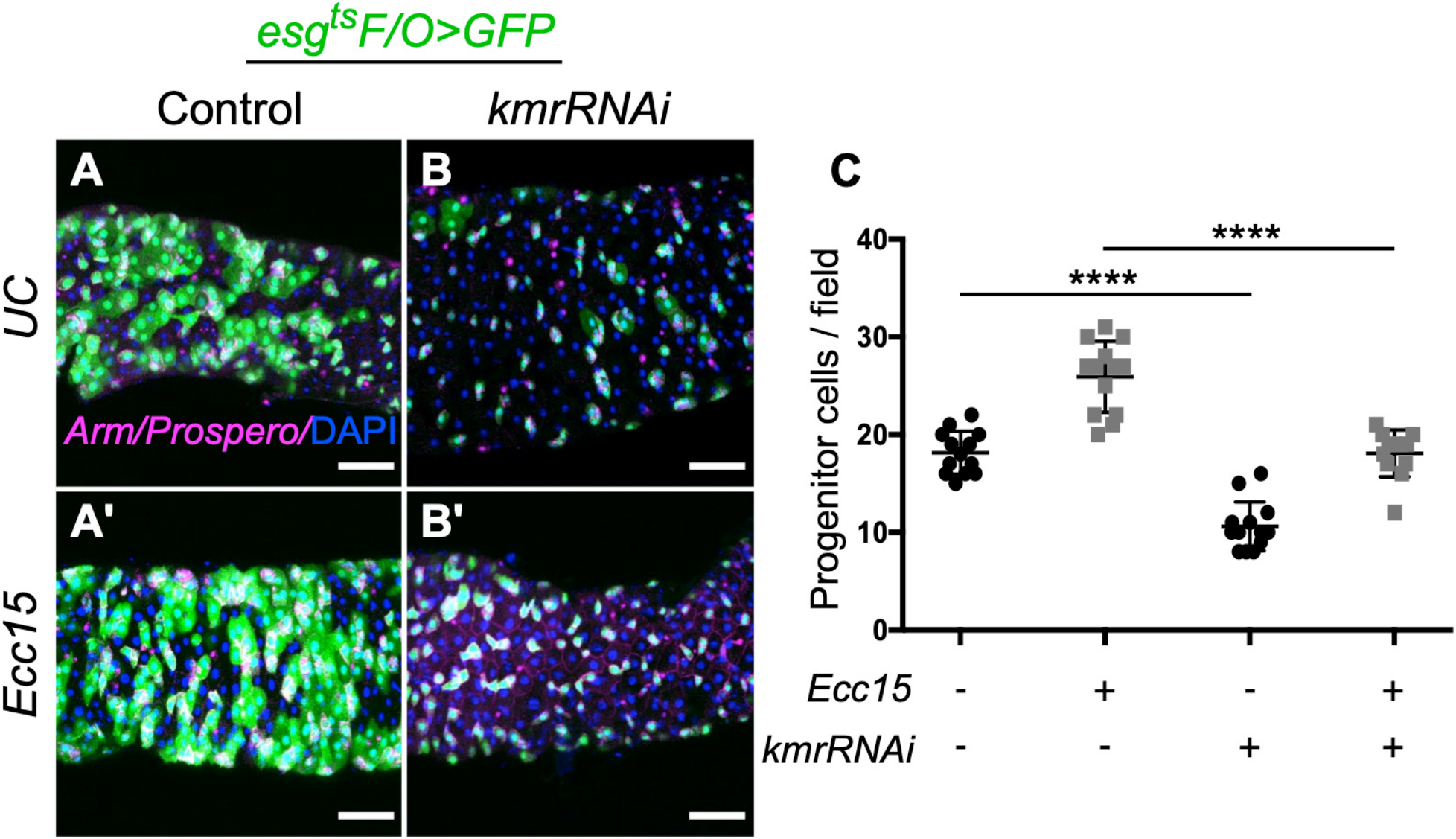
*Kramer* knockdown in intestinal stem cells and their progeny using the esg^ts^ F/O system decreases the number of progenitor cells. (A–B) Confocal micrographs of Arm/Pro staining of posterior midguts from unchallenged (UC) or *Ecc15*-challenged flies expressing *kmr* or control knockdown under the control of the *esg*^*ts*^ *F/O* system. Magenta: Arm/Pro; green: GFP (ISCs/EBs and their progeny expressing RNAi; blue: DAPI. (C) Quantification of Arm^+^ proliferating cells. One-way ANOVA (Tukey post-hoc): ****, *p*<0.0001), n=12–13. Scale bars: 40 μm.

**Figure S4 (related to Figure 5).**
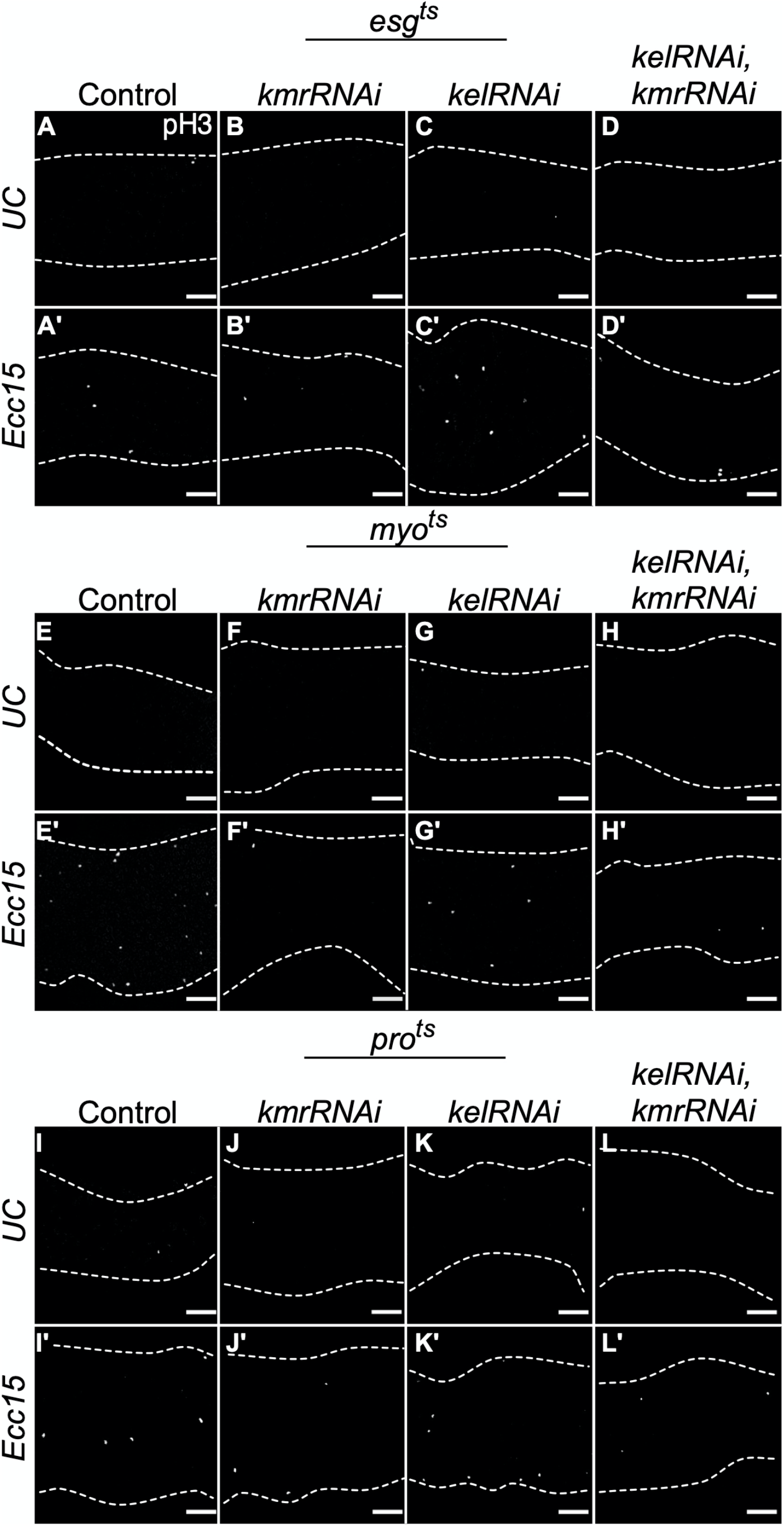
Monochrome images of pH3 staining for studies showing knockdown of *kelch* partially restores the intestinal stem cell proliferation defects caused by *kramer* downregulation in midgut cells. Shown are monochrome images of pH3 fluorescence in cells stained as described in Figure 5 (A–D, F–I, and K–N). Dashed lines indicate the boundary of the gut. Scale bars: 40 μm.

**Figure S5 (related to Figure 5).**
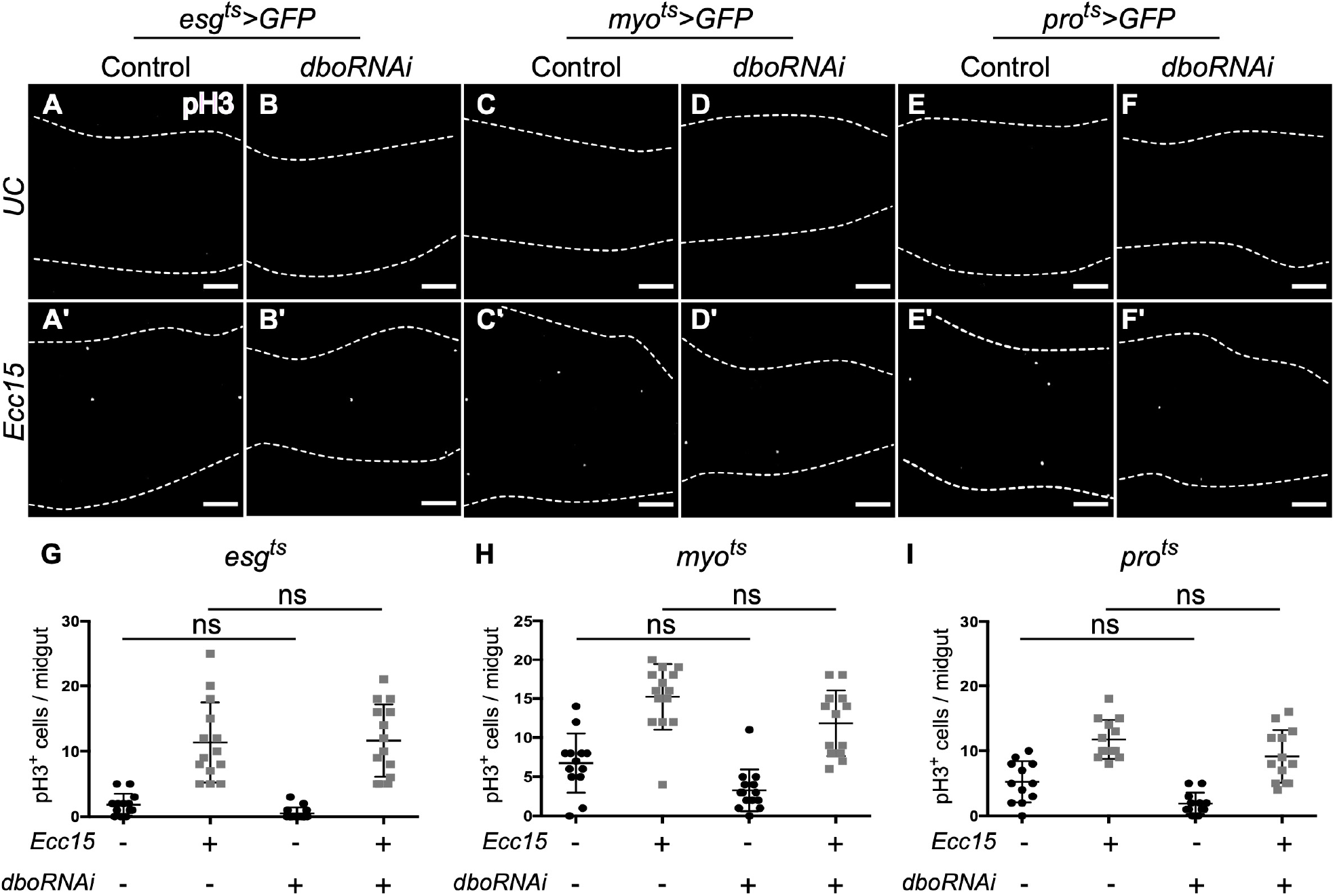
Knockdown of *diablo* (*dbo*) does not affect intestinal stem cell proliferation in the *Drosophila* midgut. Examination of proliferating cells stained with pH3 in flies expressing cell type-specific knockdown of *dbo* in ISCs/EBs (A, B, G, *esg*^*ts*^*>GFP*), ECs (C, D, H, *myo*^*ts*^*>GFP*), or EEs (E, F, I, *pro*^*ts*^*>GFP*). (A–F) Representative confocal micrographs of pH3 fluorescence. Dashed lines indicate the boundary of the gut. (G–I) Quantification of number of pH3^+^ cells. One-way ANOVA (Tukey post-hoc): ns, not significant; n=11–14. Scale bar: 40 µm.

**Figure S6 (related to Figure 5).**
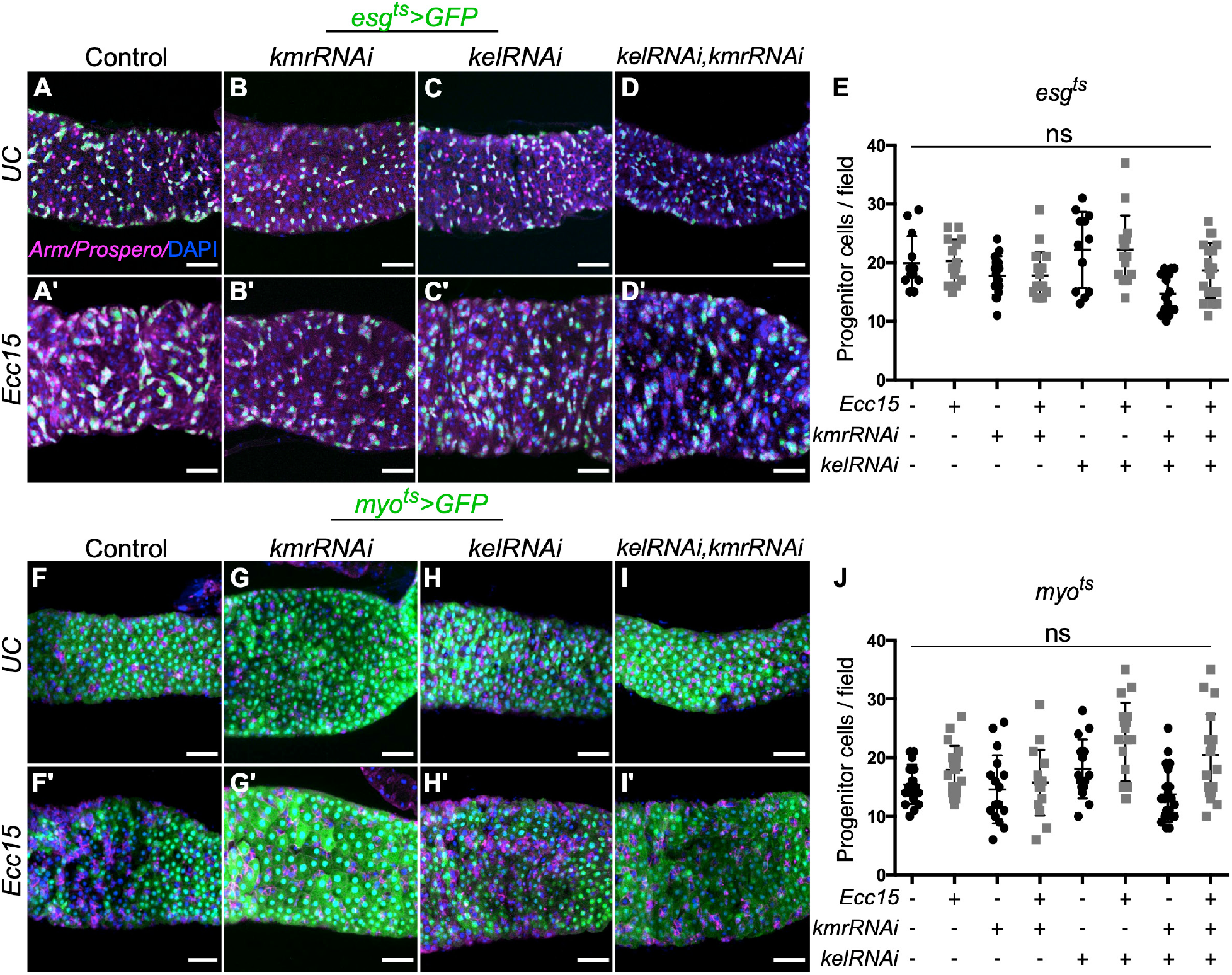
Knockdown of *kmr* and/or *kel* in ISCs/EBs or ECs has no effect on the number of Arm^+^/Pro^−^ progenitor cells in the midgut. Examination of number of Arm^+^/Pro^−^ progenitor cells in the midgut in unchallenged or *Ecc15*-challenged flies expressing conditional, cell type-specific knockdown of *kmr* and/or *kel* in ISCs/EBs (A–E, *esg*^*ts*^*>GFP*) or ECs (F–J, *myo*^*ts*^*>GFP*). Shown are representative confocal micrographs (magenta: Arm/Pro; green: GFP (RNAi)) and quantification of number of Arm^+^/Pro^−^ cells. One-way ANOVA (Tukey post-hoc): ns, not significant; n=11–14. Scale bars: 40 µm.

**Figure S7 (related to Figure 6).**
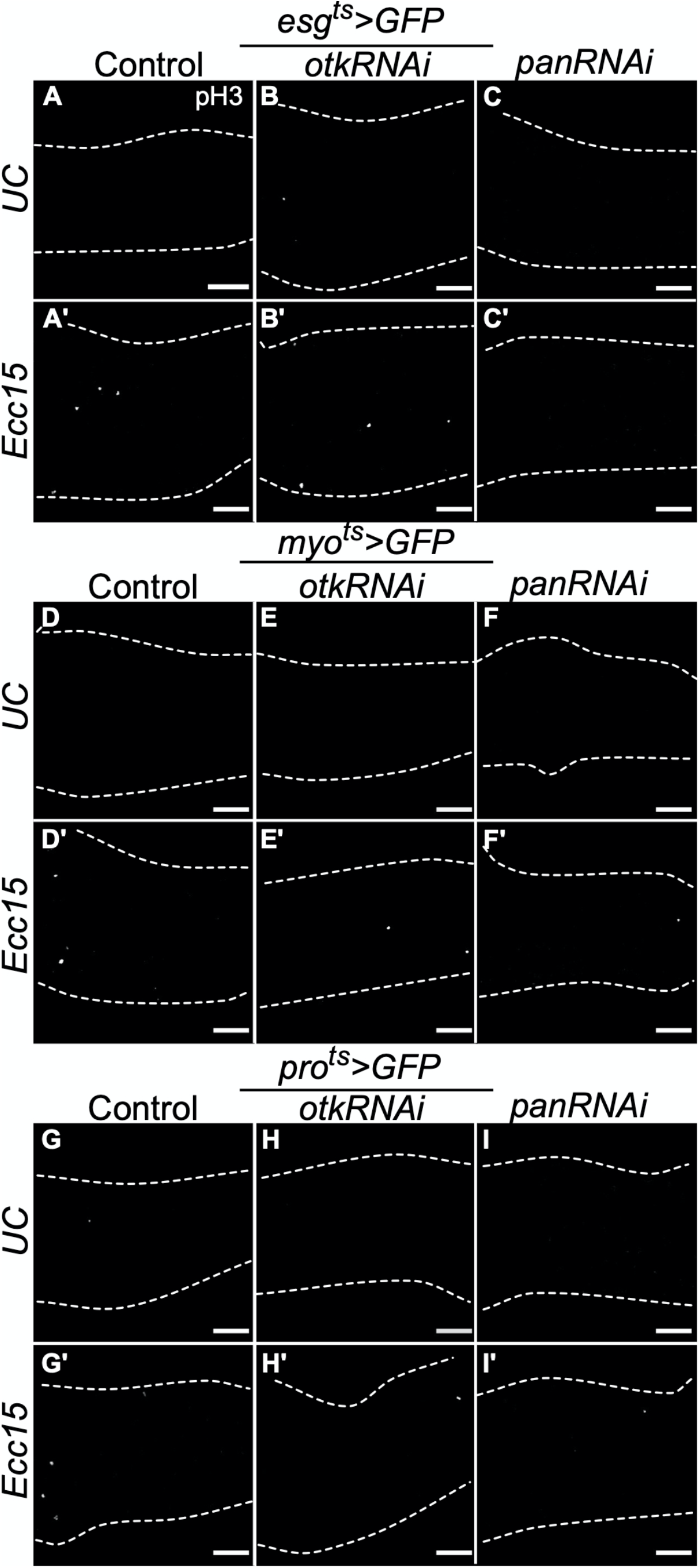
Monochrome images of pH3 staining for studies showing knockdown of *pan*, but not *otk*, phenocopies effects of *kmr* knockdown on intestinal stem cell proliferation in the midgut. Shown are monochrome images of pH3 fluorescence in cells stained as described in Figure 6 (A–C, E–G, and I–K). Dashed lines indicate the boundary of the gut. Scale bars: 40 μm.

**Table S1.**
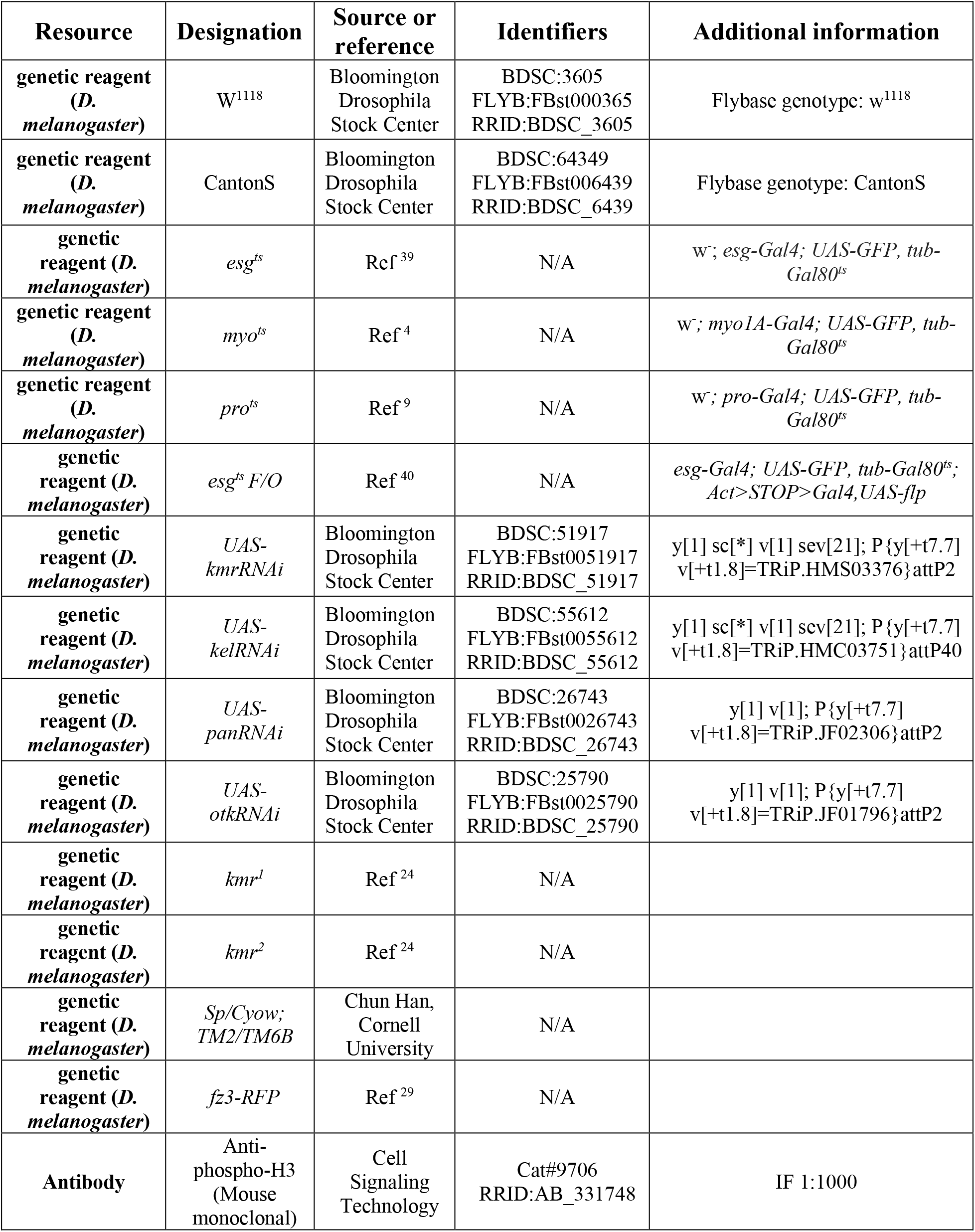
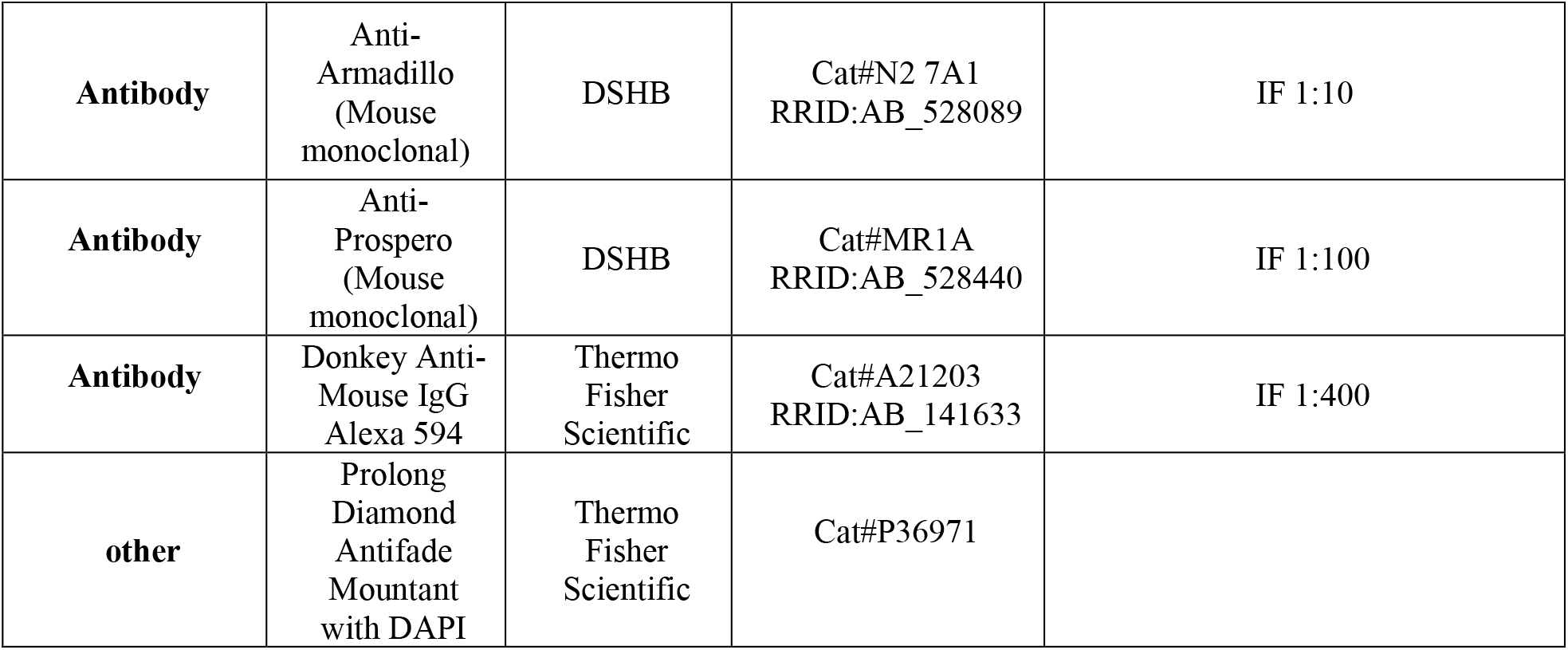
List of strains, antibodies, and reagents used in this study.

